# Thermo-Sensing Mechanisms of Splicing Control by Nuclear Stress Bodies

**DOI:** 10.1101/2025.10.21.683800

**Authors:** Tsuyoshi Ueno, Shungo Adachi, Ichiro Taniguchi, Nobuo N. Noda, Kensuke Ninomiya, Tetsuro Hirose

## Abstract

Nuclear stress bodies (nSBs) are stress-inducible membraneless organelles formed on HSATIII lncRNAs and regulate pre-mRNA splicing during thermal stress recovery. During stress, dephosphorylated SRSFs accumulate in nSBs, whereas upon stress removal, their kinase CLK1 is recruited to rephosphorylate SRSFs, thereby promoting intron detention in specific transcripts. However, the mechanism underlying CLK1 localization to nSBs in response to stress removal has remained unclear. Here, we identify phosphorylation of CLK1 at Ser341 as a critical determinant of its nSB localization. Ser341 is phosphorylated under normal conditions, dephosphorylated by PP1 during stress, and rephosphorylated by RIOK2 during recovery, enabling CLK1 localization to nSBs specifically during recovery. We further identify PPP1R2, an entirely intrinsically disordered PP1 inhibitory subunit, as a reversible thermosensor that dissociates under stress to activate PP1. Together, our findings reveal multilayered thermosensing mechanisms that coordinate the temporally staged localization of SRSFs and CLK1 to nSBs, thereby regulating temperature-dependent pre-mRNA splicing.

## INTRODUCTION

Eukaryotic cells employ diverse mechanisms to cope with environmental stresses, including transient cell cycle arrest and global reprogramming of gene expression at both transcriptional and post-transcriptional levels.^1^ Regulation of pre-mRNA splicing plays a critical role in rapid adjustment of gene expression under stresses.^2,3^ For instance, thermal stress globally suppresses pre-mRNA splicing, often through dephosphorylation of serine/arginine-rich splicing factors (SRSFs), while selectively activating the splicing of stress-response genes.^3–5^ During recovery from stresses, mechanisms that restore cellular homeostasis are activated. For example, eIF2α, a key translational inhibitor, returns to its dephosphorylated state, cancelling stress-induced translational repression.^6^ Similarly, HSP70 produced under the control of heat shock factor 1 (HSF1), binds to and inactivates HSF1, normalizing stress-induced transcription and demonstrating reversible gene expression changes.^7^

Under stress conditions, specific membraneless organelles (MLOs) are formed and served as platform of cellular responses. Cytoplasmic stress granules (SGs) and nuclear stress bodies (nSBs) are well-known examples as stress-induced MLOs.^8,9^ SGs are formed when mRNAs, untranslated due to stress, multivalently interact with RNA-binding proteins (RBPs) such as TIA-1 and G3BP1, thereby driving liquid-liquid phase separation.^10–13^ Meanwhile, nSBs are built on HSATIII long noncoding RNAs (lncRNAs),^14–16^. which belong to a class of “architectural RNAs (arcRNAs)” that serve as structural scaffolds for RBPs in the formation of MLOs.^14–17^ HSATIII arcRNAs are transcribed from primate-specific pericentromeric Satellite III (SatIII) repeat arrays located on multiple chromosomes, collectively occupying 1.5% of the human genome.^18,19^ They are enriched in GGAAU repeat sequences and exhibit heterogeneous transcript lengths.^8,20–23^ Under normal conditions, SatIII regions are heterochromatic and transcriptionally silent. Upon thermal and oxidative stress, HSF1 assembles at these regions, converting them into euchromatin and promoting transcription of HSATIII RNAs.^21,24–26^ HSATIII arcRNAs recruit a hundred of RBPs involved in pre-mRNA splicing and processing, as well as mRNA export, such as SAFB, SRSFs and SR-related proteins, to assemble nSBs.^22,26–28^ Although nSBs were first described decades ago,^29,30^ their function remained elusive until recently. We demonstrated that nSBs regulate the splicing of >400 pre-mRNAs during recovery from thermal stress through two distinct mechanisms.^22,23,31^ These mechanisms align with the general framework of MLOs, which modulate biochemical processes through three functional modes: (1) local concentration to enhance biochemical processes ("reaction crucible"), (2) sequestration of specific factors ("molecular sponge"), and (3) interactions with multiple chromatin loci ("chromatin hub").^15,32^ Within this framework, recent studies have reported that nSB formation rearranges local genes within the nSB territory to promote transcription, suggesting a role as a “chromatin hub”.^26^ In parallel, our previous work demonstrated that nSBs function as a “reaction crucible” and a “molecular sponge”, thereby modulating temperature-dependent pre-mRNA splicing.^22,31^

As the reaction crucible, nSBs facilitate temperature-dependent phosphorylation of SRSFs during stress recovery (Figure S1A).^22,23^ Under normal temperature conditions, SRSFs are highly phosphorylated, a status that is critical for their function in splicing regulation. Under thermal stress, SRSFs are globally dephosphorylated by protein phosphatase 1 (PP1), which suppresses general splicing while selectively promoting the splicing of stress-responsive pre-mRNAs.^3–5,33^ Concomitantly, HSATIII arcRNAs sequester these dephosphorylated SRSFs, to assemble nSBs. Upon return to normal temperature, CDC-like kinases (CLKs), nuclear SRSF kinases, are recruited to nSBs and rapidly rephosphorylate the associated SRSFs within 1 hour. This promotes nuclear intron retention (referred to as the “intron detention”)^34^ of target pre-mRNAs and supresses stress-induced genes. Meanwhile, as the molecular sponge, nSBs sequester N^6^-methyladenosine (m^6^A) RNA modification factors. The m^6^A writer complex components (KIAA1429 and WTAP) localize to nSBs specifically during stress recovery, resulting in m^6^A modification of adenosine in approximately 10% of HSATlll GGAAU repeat motifs. This, in turn, leads to sequestration of the m^6^A reader YTHDC1.

Consequently, nucleoplasmic availability of m^6^A modification factors is markedly reduced, limiting m^6^A modification on other pre-mRNAs. Because m^6^A facilitates efficient splicing of specific pre-mRNAs,^35^ this sequestration inhibits m^6^A-dependent splicing.^23,31^

Despite these findings, the molecular basis of temperature-dependent localization of key regulators, such as CLKs and m^6^A-related factors, to nSBs remains unclear. In this study, we focused on the crucible mechanism to elucidate how temperature controls CLK1 localization to nSBs. We show that phosphorylation of CLK1 at Ser341 promotes its localization to nSBs and enables their crucible function. Importantly, Ser341 phosphorylation is thermosensitive: it is maintained under normal conditions but lost under thermal stress. We identified RIO kinase 2 (RIOK2) as the kinase responsible for rapid rephosphorylation of Ser341 during stress recovery, whereas PP1 dephosphorylates it during thermal stress. Thus, PP1 and RIOK2 act antagonistically to regulate CLK1 localization via Ser341 phosphorylation, enabling functional switching of nSBs in response to temperature changes. Moreover, we uncovered a reversible thermosensing system in which PPP1R2-mediated PP1 activation and RIOK2-CLK1 interaction dynamically modulate CLK1 phosphorylation and localization, coordinating nSB function for rapid adaptation to temperature shifts.

## RESULTS

### Three distinct regions within CLK1 IDR are essential for its temperature-dependent localization to nSBs

To elucidate the molecular mechanism underlying the temperature-dependent CLK1 localization to nSBs as a key step of their molecular crucible function (Figure S1A), we first aimed to identify the essential regions within CLK1 required for this process. In this study, to simplify the validation of CLK1 localization to nSBs by excluding potential effects of substrate phosphorylation within nSBs, we introduced a kinase-dead mutation (K191R) for the localization analysis (Figure S1B). The CLK1 K191R mutant maintains temperature-dependent localization to nSBs: it does not localize to nSBs under thermal stress, but appropriately localizes to nSBs during stress recovery (Figure S1C). This enables us to investigate the localization mechanism independently of CLK1 kinase activity. Our previous study demonstrated that the N-terminal IDR of CLK1 (Figure S1D) is both necessary and sufficient for its localization to nSBs.^22^ Specifically, the IDR-deletion mutant (ΔIDR) fails to localize to nSBs, whereas the CLK1 IDR alone is capable of temperature-dependent localization to nSBs (Figures S1E-S1H).

To gain further mechanistic insights into CLK1 localization to nSBs via its IDR, we generated a series of CLK1 IDR partial deletion mutants. To design the partial deletion constructs, we referred to CLK2 and CLK4, which are the other members of CLK family and are also recruited to nSBs through their IDRs.^22^ While the kinase domain is highly conserved among CLK family proteins (Figure S1I), their IDRs are only partially conserved, with several conserved regions within these IDRs (Figure S1J). Based on this observation, we divided the CLK1 IDR into five segments and generated deletion mutants lacking each segment (Δ1- Δ5) (Figures 1A, S1J, and S1K). The localization of these mutants to nSBs was examined by RNA fluorescence in situ hybridization (RNA-FISH) of HSATIII combined with immunofluorescence (IF) of FLAG-CLK1 and quantified by calculating the correlation coefficient between the HSATIII and FLAG-CLK1 signals in the nucleus. The nSB localization of the Δ1, Δ3, and Δ5 CLK1 mutants was significantly reduced (Figures 1B and 1C), indicating that these three regions are required for CLK1 localization to nSBs. Based on these findings, we constructed Δ1,3,5 mutant lacking 1, 3, and 5, and Δ2,4 mutant retaining these regions (Figures 1A and S1L). The Δ1,3,5 mutant failed to localize to nSBs, whereas the Δ2,4 mutant localized to nSBs similarly to the full-length (FL) CLK1 (Figures 1D and 1E). To corroborate these findings, we conducted HSATIII-ChIRP (chromatin isolation by RNA purification) followed by western blotting (WB), which purifies HSATIII-RNP complexes using a biotinylated antisense oligonucleotide (ASO) to detect the copurified CLK1 variants (Figure 1F). The pulldown efficiency of the Δ1,3,5 mutant during thermal stress recovery was significantly lower than those of the FL and Δ2,4 mutants (42□→37□ in Figures 1G and 1H). None of the CLK1 variants were precipitated during the thermal stress (42□ in Figures 1G and 1H). These data suggest that three distinct regions within CLK1 IDR are essential for its temperature-dependent localization to nSBs during stress recovery. We named these regions “nSB <u>L</u>ocalization <u>R</u>egions (nSB-LRs)” (Figure 1A).

**Figure 1.**
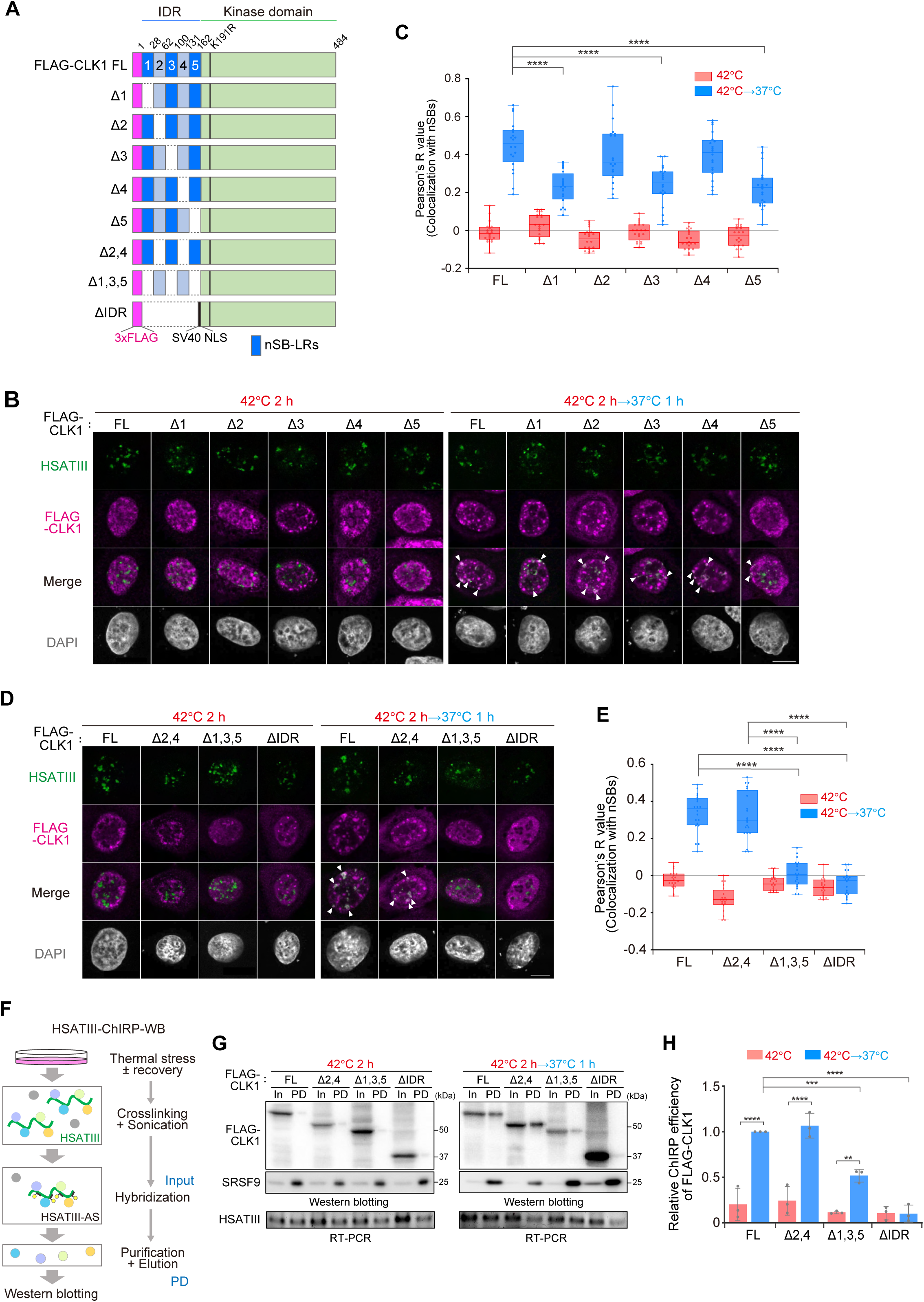
Three distinct regions within CLK1’s IDR are essential for its temperature-dependent localization to nSBs. (A) Schematic of domain structures of CLK1 mutants and nSB-LRs. (B, D) Subnuclear localization of CLK1 mutants under thermal stress and recovery, visualized by HSATIII RNA-FISH (green) combined with FLAG immunofluorescence (IF) (magenta). White arrowheads indicate HSATIII-CLK1 colocalization. Nuclei were counterstained with DAPI. Scale bar, 10 μm. (C, E) Quantification of colocalization between CLK1 mutants and nSBs in (B, D). Box plots show Pearson’s correlation coefficients (R values) between HSATIII-FISH and FLAG IF signals in the nucleus (n = 20). \*\*\*\**p* < 0.0001 (Dunn’s multiple comparisons test). (F) Overview of the HSATIII-ChIRP assay using biotinylated ASO targeting HSATIII. (G) Analysis of HSATIII-associated FLAG-CLK1 and endogenous SRSF9 by ChIRP-western blotting (WB). Protein input (In) 5%. HSATIII lncRNA was detected by RT-PCR, with 25% of the total RNA input loaded (In). PD, pulldown products of HSATIII-ChIRP. (H) Quantification of ChIRP efficiency from (G). ChIRP efficiency was determined by calculating the ratio of PD to In. Data are presented as mean ± SD (n = 3). \*\*\*\**p* < 0.0001, ***0.0001 < *p* < 0.001, **0.001 < *p* <0.01 (Šidák’s multiple comparisons test).

### RIOK2 mediates temperature-dependent localization of CLK1 to nSBs

We hypothesized that the regulatory proteins involved in CLK1 localization to nSBs bind to CLK1 via nSB-LRs. To identify them, we performed a coimmunoprecipitation–mass spectrometry (coIP-MS)–based screen using both full-length CLK1 and deletion mutants with or without nSB-LRs. By comparing their binding partners, we identified several candidate regulators that specifically bind to nSB-LRs (Table S1). We first validated their interactions by coIP followed by western blotting (coIP-WB), and confirmed that four candidates (RIOK2, NAGK, AHCYL1, and ERP44) bind to CLK1 in an nSB-LR-dependent manner (Figure 2A). To assess their contribution to CLK1 localization to nSBs, we knocked down each candidate and analyzed CLK1 localization (Figures 2B-2D and S2A-S2F). Knockdown of the serine/threonine kinase RIO kinase 2 (RIOK2) significantly inhibited CLK1 localization to nSBs (Figures 2B-2D). Consistently, HSATIII-ChIRP revealed reduced association of CLK1 with HSATIII arcRNAs upon RIOK2 knockdown (Figures S2G and S2H). In contrast, knockdown of the other regulator candidates had no apparent effect on CLK1 localization (Figures S2A-S2F). Based on these findings, we focused on RIOK2 as a key regulator of CLK1 localization to nSBs. RIOK2 functions in both the nucleus and the cytoplasm; it is required for cytoplasmic 40S ribosomal subunit maturation^36,37^ and acts as a nuclear transcriptional activator of hematopoietic transcription factors involved in human blood cell differentiation.^38^ Consistently, our IF analysis showed that RIOK2 is mainly distributed in the cytoplasm, with a minor population also detectable in the nucleus (Figures S2I-S2K). Furthermore, HSATIII-FISH revealed that RIOK2 is poorly localized to nSBs during and after thermal stress (Figure S2I).

**Figure 2.**
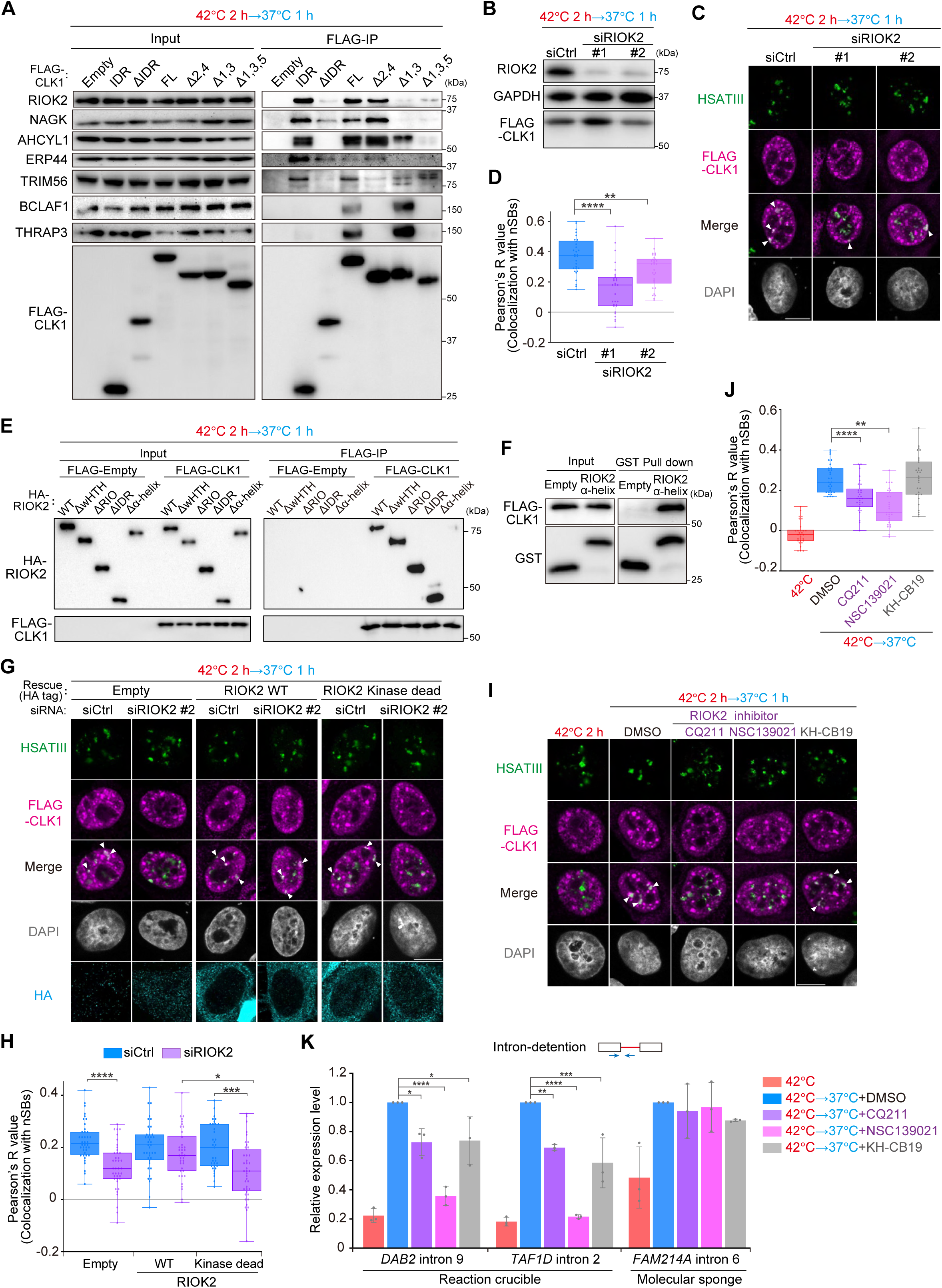
RIOK2 regulates temperature-dependent localization of CLK1 to nSBs. (A) CLK1 coIP from cell lysates during stress recovery, followed by WB. FLAG-CLK1 variants used as bait for coIP are indicated above the panels. (B) Evaluation of RIOK2 knockdown efficiency by WB. siRNAs (20 nM) are indicated above the upper panel. GAPDH is a loading control. (C, G, I) Subnuclear localization of CLK1 under thermal stress and recovery, visualized by HSATIII RNA-FISH combined with FLAG IF. (C, G) Cells were transfected with siRNAs (20 nM). (I) DMSO or RIOK2 inhibitors (CQ211, 10 μM; NSC139021, 30 μM) were added during recovery. Scale bar, 10 μm. (D, H, J) Quantification of colocalization between CLK1 and nSBs (n = 30) in (C, G, I). \*\*\*\**p* < 0.0001, ***0.0001 < *p* < 0.001, **0.001 < *p* <0.01, *0.01 < *p* < 0.05 (Dunn’s multiple comparisons test). (E) CoIP of CLK1 with RIOK2 mutants from cell lysates during stress recovery, followed by WB. HA-RIOK2 variants analyzed by coIP are indicated above the panels. (F) Analysis of direct interactions between recombinant CLK1 and isolated α-helical fragments of RIOK2 by GST pulldown and WB. (K) Monitoring of intron-retaining pre-mRNA levels during stress and recovery by qRT-PCR. Data are presented as mean ± SD (n = 3). Pre-mRNA levels were calculated as the ratio of each RNA to *18S* rRNA and were normalized to the 42°C→37°C+DMSO condition. DMSO or RIOK2 inhibitors (CQ211, 10 μM; NSC139021, 30 μM) were added during recovery. \*\*\*\**p* < 0.0001, ***0.0001 < *p* < 0.001, **0.001 < *p* <0.01, *0.01 < *p* < 0.05 (Dunnet’s multiple comparisons test).

To gain mechanistic insight into the RIOK2-CLK1 interaction, we sought to identify the region within RIOK2 that is required for this interaction. RIOK2 consists of a DNA-binding winged helix-turn-helix (wHTH) domain, a kinase (RIO) domain, an IDR, and a C-terminal α-helix (Figure S2L).^38,39^ We generated RIOK2 deletion mutants lacking each domain (Figure S2M) and examined their binding to CLK1 using coIP-WB. This analysis revealed that deletion of the C-terminal α-helix markedly impaired RIOK2-CLK1 interaction (Figure 2E). To corroborate these findings, we purified recombinant proteins comprising the RIOK2 α-helix alone fused to GST- and His-tag, together with FLAG-tagged CLK1 (Figure S2N). An in vitro binding assay revealed that the isolated α-helix alone was sufficient for binding to CLK1 (Figure 2F). These results demonstrate that RIOK2 directly binds to nSB-LRs of CLK1 via its C-terminal α-helix.

To evaluate the contribution of RIOK2 kinase activity to CLK1 localization, we performed a knockdown and rescue experiment using a wild-type RIOK2 or a kinase-dead RIOK2 mutant (K123A, D246A)^40^ (Figure S2O). While wild-type (WT) RIOK2 successfully rescued CLK1 localization to nSBs under RIOK2 knockdown conditions (siRIOK2 #2), the kinase-dead mutant did not (Figures 2G and 2H). To inhibit RIOK2 kinase activity specifically during stress recovery, we added RIOK2 chemical inhibitors to the cells after thermal stress removal and evaluated CLK1 localization. The RIOK2 inhibitors CQ211^41^ and NSC139021^42^ significantly inhibited CLK1 localization to nSBs, whereas the CLK1 inhibitor KH-CB19^43^ had no effect (Figures 2I and 2J). These data indicate that RIOK2 kinase activity is essential for proper CLK1 localization to nSBs. CLK1 also localizes to nuclear speckles regardless of thermal conditions. Nuclear speckles are prominent nuclear compartments enriched in SRSFs and play key roles in pre-mRNA splicing.^22^ Interestingly, neither RIOK2 knockdown nor RIOK2 inhibitors affected CLK1 localization to nuclear speckles (Figures S2P-S2R), indicating that RIOK2 selectively regulates CLK1 localization to nSBs.

Next, we examined the hypothesis that RIOK2 facilitates nSB-mediated pre-mRNA splicing by promoting CLK1 localization. We previously reported that nSBs promote target intron detention during stress recovery through two distinct mechanisms: (1) the reaction crucible mechanism, in which CLK1 rephosphorylates SRSFs and SR-related proteins, and (2) the molecular sponge mechanism, which involves sequestration of m^6^A modification factors.^22,23,31^ These mechanisms predominantly affect distinct subsets of pre-mRNAs. Thus, if our hypothesis is correct, RIOK2 inhibition should specifically impair the reaction-crucible-dependent splicing. Indeed, RIOK2 inhibition during stress recovery significantly suppressed intron detention of *DAB2* and *TAF1D* mRNAs, which depend on the reaction crucible, similarly to CLK1 inhibition as a positive control. In contrast, intron detention of *FAM214A*, which depends on the molecular sponge, was unaffected by RIOK2 inhibition (Figure 2K). These findings indicate that RIOK2 facilitates CLK1 localization to nSBs, thereby specifically promoting temperature-dependent splicing regulation through the nSB crucible mechanism.

### RIOK2-dependent phosphorylation of CLK1 at Ser341 drives its nSB localization

Next, we aimed to elucidate the molecular mechanism by which RIOK2 facilitates CLK1 localization to nSBs during stress recovery. Given that RIOK2 kinase activity is required for proper CLK1 localization to nSBs (Figures 2G-2J), we hypothesized that RIOK2 phosphorylates CLK1. To verify this, we performed Phos-tag™ SDS-PAGE followed by WB to analyze the phosphorylation status of CLK1. As shown in Figure 3A, we found that CLK1 is phosphorylated at normal temperature condition (37□) and transiently dephosphorylated during thermal stress (42□ 2 h) and rephosphorylated during stress recovery (42 J 2 h→37 1 h). Notably, treatment with RIOK2 inhibitors during stress recovery prevented CLK1 rephosphorylation, indicating that RIOK2 mediates this modification (Figure 3A). To identify RIOK2-dependent phosphorylation sites within CLK1, we analyzed phosphorylated peptides by MS and found that 323-343 amino acid (aa) region of CLK1 is phosphorylated by RIOK2 in a temperature-dependent manner (Figure 3B). We also confirmed that the GAPDH Ser83 phosphorylation, used as a control, was not significantly affected either by temperature changes or RIOK2 inhibition, indicating that the observed changes in CLK1 phosphorylation do not result from biases in total protein input after phosphopeptide purification (Figure S3A). Based on PhosphoSitePlus^®^ database, 323-343 aa region of CLK1 contains four serine/threonine residues that are potential phosphorylation sites (Figure S3B). We generated alanine substitution mutants of each site using the kinase-dead CLK1 mutant (K191R refer to as the "control”). In addition, we created a combined mutant in which all serine/threonine residues in this region were substituted with alanine (referred to as the S/T→A mutant) (Figure S3C). Localization analysis revealed that the S341A mutant failed to localize to nSBs, similar to the S/T→A mutant (Figures 3C and 3D). Moreover, the S341A mutant mislocalized from nSBs in a manner similar to the control CLK1 treated with RIOK2 inhibitor (Figures 3E and 3F). Importantly, treatment with the RIOK2 inhibitor had no additional effect on the S341A mutant (Figures 3E and 3F), suggesting that CLK1 Ser341 is a RIOK2-dependent phosphorylation site that is essential for its proper localization to nSBs. To verify this, we performed an in vitro kinase assay using purified recombinant RIOK2 and CLK1 proteins (Figure S3D). In this system, phosphorylation of the CLK1 protein depended on RIOK2 (Figure S3E), and we confirmed that phosphorylation of both RIOK2 and CLK1 was inhibited by the RIOK2 inhibitor in a dose-dependent manner (Figure S3F). Phosphorylation of the CLK1 S341A mutant was significantly reduced compared with the control CLK1, indicating that RIOK2 directly phosphorylates CLK1 at Ser341 (Figures 3G and 3H). Based on these results, we conclude that RIOK2 phosphorylates CLK1 at Ser341 during stress recovery, facilitating the temperature-dependent localization of CLK1 to nSBs.

**Figure 3.**
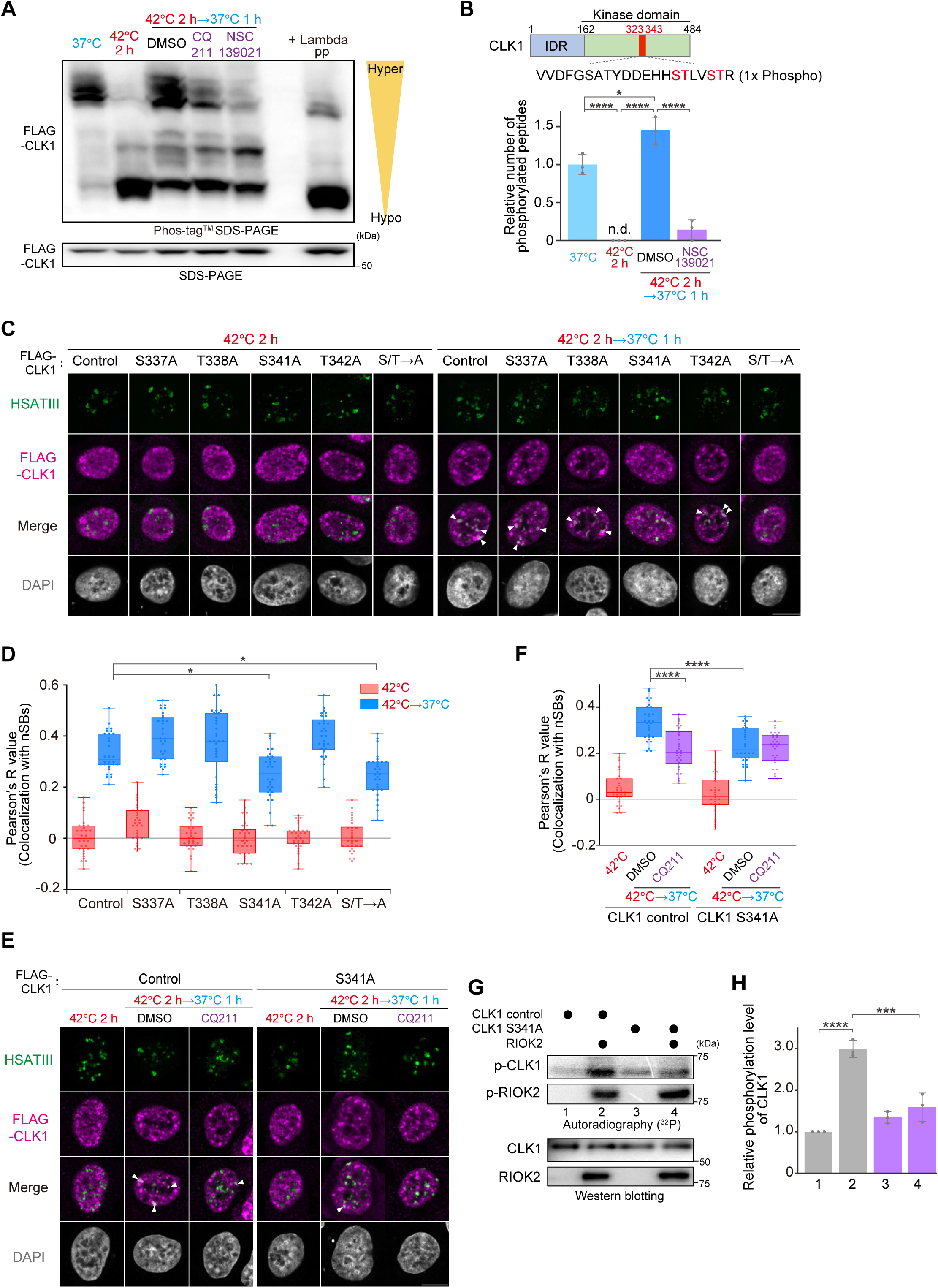
RIOK2 phosphorylates CLK1 at Ser341 to promote its localization to nSBs and splicing regulation via nSBs. (A) Phos-tag™ SDS-PAGE (upper panel) and standard SDS-PAGE (lower panel) followed by WB of CLK1. Migration of different phosphorylation states is shown on the right. Lambda phosphatase treated sample (+Lambda pp) was used as a reference for dephosphorylated CLK1. DMSO or RIOK2 inhibitors (CQ211, 20 μM; NSC139021, 30 μM) were added during recovery. (B) Schematic of the CLK1 peptide spanning amino acids (aa) 323–343 and bar graph showing the abundance of the phosphorylated peptide across temperature conditions. DMSO or RIOK2 inhibitors were added during recovery as in (A). “1x Phospho” denotes phosphorylation at a single Ser/Thr/Tyr residue. n.d., below the detection limit. Data are presented as mean ± SD (n = 3), with each sample measured twice by mass spectrometry and averaged. \*\*\*\**p* < 0.0001, *0.01 < *p* < 0.05 (Šidák’s multiple comparisons test). (C, E) Subnuclear localization of CLK1 mutants under thermal stress and recovery, visualized by HSATIII RNA-FISH combined with FLAG IF. (C) S/T→A mutant carries alanine substitutions at all serine/threonine residues within CLK1 aa 323-343 (S328A/S330A/S337A/T338A/S341A/T342A). (E) DMSO or CQ211 (10 μM) was added during recovery. Scale bar, 10 μm. (D, F) Quantification of colocalization between CLK1 mutants and nSBs (n = 30) in (C, E). \*\*\*\**p* < 0.0001, *0.01 < *p* < 0.05 (Dunn’s multiple comparisons test). (G) In vitro phosphorylation of RIOK2 and CLK1 assessed by autoradiography with WB validation. (H) Quantification of autoradiographic signal intensities from (G). Each bar corresponds to the lane numbers shown in (G). Data are presented as mean ± SD (n = 3). \*\*\*\**p* < 0.0001, ***0.0001 < *p* < 0.001 (Šidák’s multiple comparisons test).

Additionally, to assess whether the phosphorylation at S341 affects CLK1 kinase activity, we purified enzymatically active CLK1 WT and S341A mutant (Figure S3G). An in vitro kinase assay revealed that the autophosphorylation activity of the S341A mutant was slightly weaker than that of CLK1 WT (Figures S3H and S3I), suggesting that Ser341 phosphorylation by RIOK2 activates CLK1. CLK1 kinase activity is essential for splicing regulation by nSBs, as shown by inhibition of nSB-mediated splicing upon treatment with KH-CB19, a CLK1 inhibitor (Figure 2K). Therefore, RIOK2-mediated phosphorylation of CLK1 may promote CLK1-dependent SRSF phosphorylation and subsequent splicing regulation via nSBs.

The CLK family consists of CLK1-4. Among them, CLK1, CLK2, and CLK4 have been shown to localize to nSBs.^22^ We aligned the amino acid sequences of CLK1-4 surrounding the Ser341 region of CLK1 and found that this serine is uniquely substituted with alanine only in CLK3 (Figure S3J). We constructed plasmids expressing each CLK paralogue (Figure S3K) and evaluated their localization to nSBs as well as the role of RIOK2 in this process. As a result, CLK1, CLK2, and CLK4 localized to nSBs during thermal stress recovery in a RIOK2-dependent manner, whereas CLK3 did not localize to nSBs and remained unaffected by RIOK2 inhibition (Figures S3L and S3M). These findings further support that RIOK2 selectively contributes to the localization of CLKs to nSBs, correlating with the presence of a specific phosphorylation site.

### PP1 dephosphorylates CLK1 at Ser341 during thermal stress

As described above, RIOK2 binds to CLK1 via its nSB-LRs and phosphorylates Ser341 during stress recovery, thereby facilitating CLK1 localization to nSBs and the subsequent splicing regulation by nSBs. The phosphorylation state at this site is thus crucial for the functional switching of nSBs. We therefore sought to identify the phosphatase responsible for dephosphorylation of CLK1 at Ser341 during thermal stress. In our CLK1 coIP-MS analysis, we identified six candidate protein phosphatases that bind to CLK1 (Figure 4A). Among these, all three catalytic subunits of PP1 (PPP1CA, PPP1CB, and PPP1CC) were identified, and notably, they were the only phosphatases localized exclusively in the nucleus. Therefore, we focused on PP1 as a candidate phosphatase that dephosphorylates CLK1 during thermal stress. Because PPP1CC exhibited the strongest binding to CLK1 among the three catalytic subunits, we proceeded with further analysis using PPP1CC. CoIP analysis confirmed that PPP1CC interacts with CLK1 in cells (Figure 4B). Although PP1 catalytic subunits associate with the IDR of CLK1, they did not display a clear region-specific interaction, in contrast to RIOK2 (Table S1 and Figure S4A). Moreover, Phos-tag WB revealed that overexpression (OE) of PPP1CC, but not of the PP2A catalytic subunit (PPP2CA), promoted CLK1 dephosphorylation, suggesting that PP1 dephosphorylates CLK1 in cells (Figures 4C and S4B).

**Figure 4.**
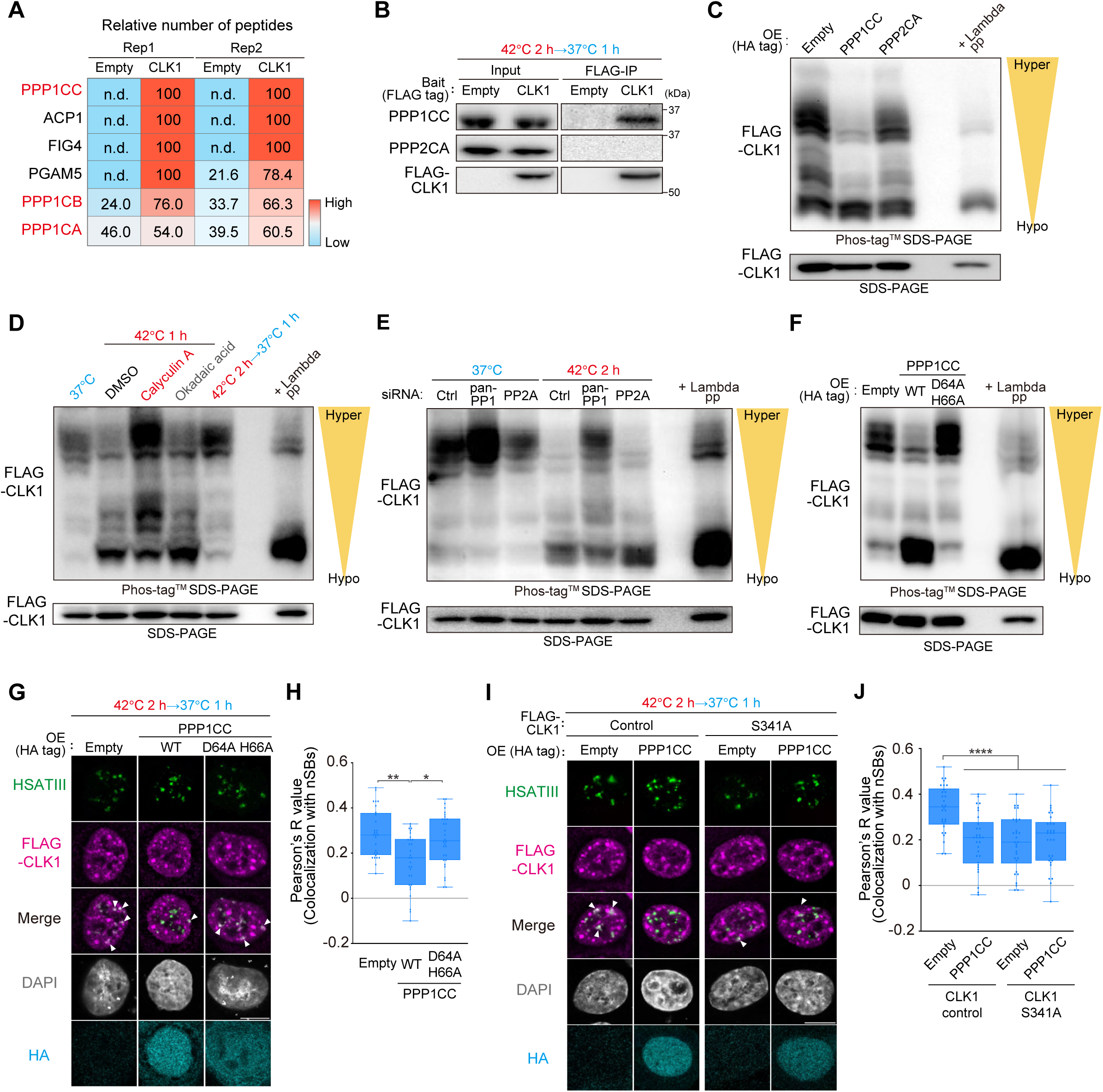
PP1 dephosphorylates CLK1 at Ser341 during thermal stress. (A) Heat map of peptides identified by mass spectrometry following coIP from cell lysates during stress recovery, with either vector control (Empty) or CLK1. Data are shown as relative number of peptides coimmunoprecipitated, averaged from two MS measurements for each sample. (B) CoIP from cell lysates during stress recovery and WB analysis of CLK1 binding to PPP1CC and PPP2CA. (C–F) Phosphorylation status of FLAG-CLK1 analyzed by Phos-tag™ SDS-PAGE (upper panel) and standard SDS-PAGE (lower panel), followed by WB. (D) Effect of phosphatase inhibitors (Calyculin A, 5 nM; Okadaic acid, 2 nM) or DMSO during thermal stress. (E) Effect of PP1 or PP2A knockdown (20 nM). (F) Effect of overexpression (OE) of WT PPP1CC or the catalytically inactive D64A/H66A mutant. (G, I) Subnuclear localization of CLK1 (control and mutants) in PPP1CC (WT and mutant) OE cells during recovery, visualized by HSATIII RNA-FISH combined with FLAG IF. Scale bar, 10 μm. (H, J) Quantification of CLK1 colocalization with nSBs (n = 30) from (G, I). \*\*\*\**p* < 0.0001, **0.001 < *p* <0.01, *0.01 < *p* < 0.05 (Dunn’s multiple comparisons test).

To further assess whether PP1 dephosphorylates CLK1 during thermal stress, we examined the effects of phosphatase inhibitors on CLK1 phosphorylation. Okadaic acid was employed to selectively inhibit PP2A, while calyculin A was used to inhibit both PP1 and PP2A. Consistent with a role for PP1 in SRSF dephosphorylation,^33^ WB with a pan-phospho-SR antibody (1H4) showed that thermal stress-induced dephosphorylation of SRSFs was blocked by calyculin A but not by okadaic acid (Figure S4C). Similarly, Phos-tag WB revealed that CLK1 dephosphorylation during thermal stress was inhibited by calyculin A but not by Okadaic acid (Figure 4D). Consistently, PP1 knockdown impaired CLK1 dephosphorylation under thermal stress, whereas PP2A knockdown had no effect (Figures 4E and S4D). Together, these data indicate that PP1 mediates CLK1 dephosphorylation in response to thermal stress.

Next, we assessed whether PP1 directly dephosphorylates CLK1 at Ser341. If so, PP1 OE should inhibit CLK1 localization to nSBs and have no additional effect on the S341A mutant. As expected, PPP1CC OE inhibited CLK1 localization to nSBs, whereas a catalytically inactive PPP1CC mutant (D64A, H66A)^44,45^ did not (Figures 4F-4H and S4E). Furthermore, PPP1CC OE had no additional effect on the S341A mutant (Figures 4I, 4J, and S4F), although PPP1CC still bound to the mutant (Figure S4G). These findings suggest that PPP1CC OE prevents RIOK2-dependent CLK1 localization to nSBs and that PP1 dephosphorylates CLK1 at Ser341. Based on these results, we conclude that RIOK2 and PP1 function in opposition to regulate the phosphorylation status of CLK1, thereby controlling temperature-dependent functional switching of nSBs.

### RIOK2 preloads on CLK1 during thermal stress

We further analyzed the molecular basis of the thermosensing mechanisms of RIOK2 and PP1, which are crucial for temperature-dependent functional switching of nSBs. First, we confirmed that the expression levels and intracellular localization of RIOK2 and PP1 remained largely unchanged in response to temperature shifts (Figures S2I, S5A, and S5B). Neither RIOK2 nor PP1 was detectable in nSBs under any conditions. We then examined whether temperature affects the interactions between CLK1 and RIOK2 or between CLK1 and PP1. Interestingly, the interaction levels of both CLK1-RIOK2 and CLK1-PP1 were significantly increased under thermal stress (42□ in Figures 5A and 5B) compared with normal (37□) and recovery (42□→37□) conditions. Similar results were obtained using EGFP-tagged CLK1 coIP, confirming the finding from FLAG-tagged CLK1 coIP (Figures S5C and S5D).

**Figure 5.**
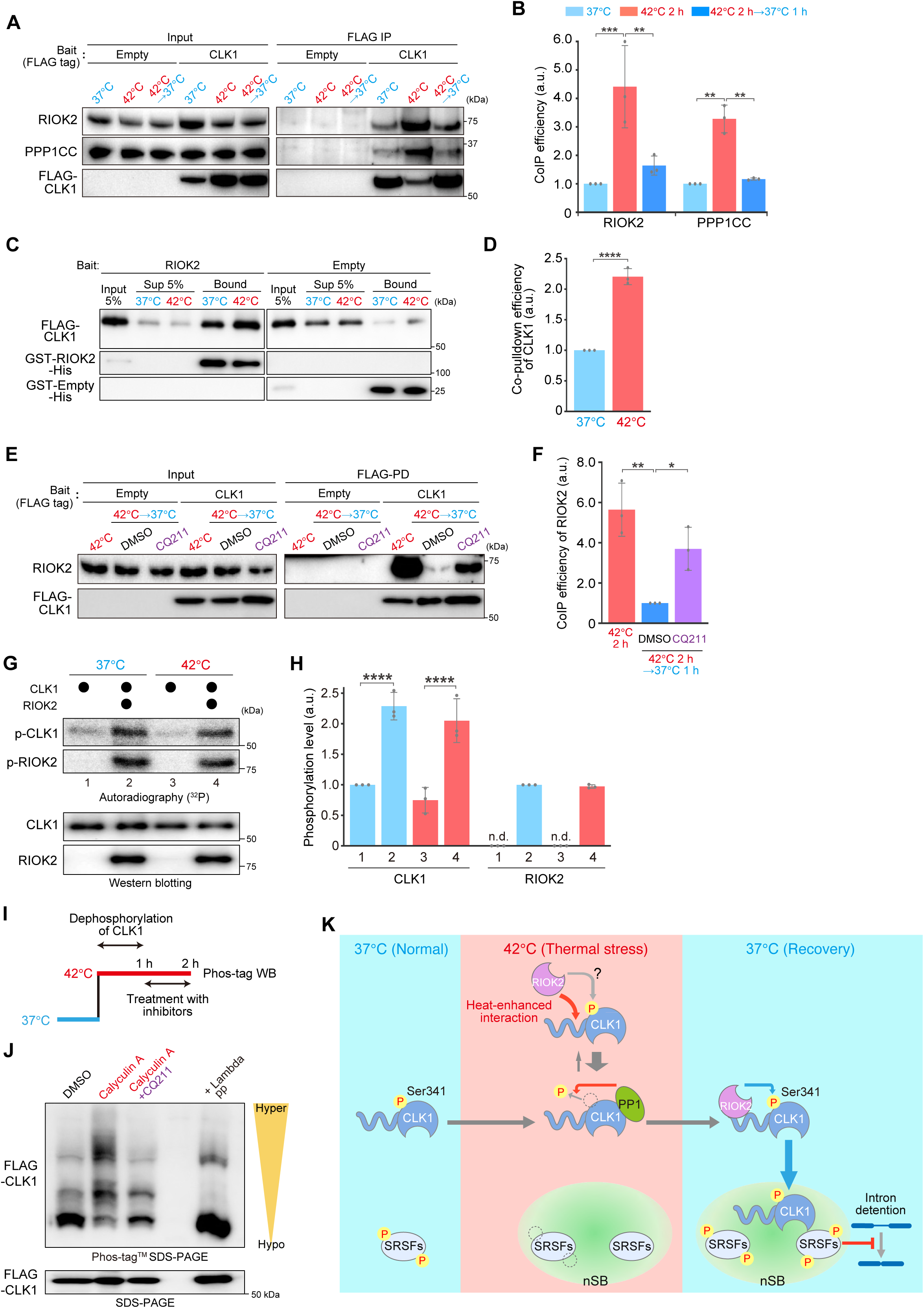
RIOK2 is preloaded onto CLK1 during thermal stress. (A) CLK1 interactions with RIOK2 and PPP1CC across temperature conditions analyzed by FLAG-CLK1 coIP and WB to detect RIOK2 and PPP1CC. (B, F) Quantification of CoIP in (A,E). CoIP efficiency was calculated as the ratio of FLAG IP band intensity of RIOK2 or PPP1CC to that of CLK1. Data are presented as mean ± SD (n = 3). Statistical significance was determined by Šidák’s (B) or Tukey’s (F) multiple comparisons test. (***0.0001 < *p* < 0.001, **0.001 < *p* <0.01, *0.01 < *p* < 0.05). (C) Direct interaction between recombinant CLK1 and RIOK2 proteins at 37□ or 42□ assessed by GST pulldown of RIOK2 and WB to detect FALG-CLK1. (D) Quantification of co-pulldown efficiency of CLK1 in (C). Co-pulldown efficiency was calculated as the ratio of the band intensity of CLK1 to that of RIOK2 in the bound fraction. Data are presented as mean ± SD (n = 3). \*\*\*\**p* < 0.0001 (two-tailed t-test). (E) Effect of RIOK2 inhibitors on CLK1-RIOK2 interactions during stress recovery. Lysates were prepared from cells exposed to thermal stress alone or from cells recovering from thermal stress in the presence of DMSO or CQ211 (10 μM) and subjected to FLAG-CLK1 coIP followed by WB for RIOK2. (G) In vitro phosphorylation of RIOK2 and CLK1 at 37□ or 42□ analyzed by autoradiography with western blot validation. (H) Quantification of autoradiographic signals in (G). Each bar corresponds to the lane numbers shown in (G). n.d., below the detection limit. Data are presented as mean ± SD (n = 3). ***0.0001 < *p* < 0.001 (Šidák’s multiple comparisons test). (I) Schematic of the experimental design to assess RIOK2-mediated phosphorylation of CLK1 under thermal stress. (J) Analysis of RIOK2-mediated phosphorylation of CLK1 during stress. Phosphorylation status of FLAG-CLK1 was analyzed by Phos-tag™ SDS-PAGE (upper panel) and standard SDS-PAGE (lower panel), followed by WB. After 1 hour of thermal stress, DMSO or inhibitors (Calyculin A, 5 nM; CQ211, 20 nM) were added and thermal stress was continued, as indicated (I).

We initially focused on the enhanced binding of RIOK2 to CLK1 under thermal stress and considered two possibilities: (1) RIOK2 preferentially binds to dephosphorylated CLK1 produced by PP1 during thermal stress, or (2) the binding of RIOK2 to CLK1 is intrinsically enhanced at 42□, independent of CLK1’s phosphorylation status. To address the first possibility, we overexpressed PP1 and performed coIP-WB. The binding of RIOK2 to CLK1 was not enhanced by PP1 OE (Figures S5E and S5F). To further investigate this, recombinant GST-RIOK2-His and FLAG-CLK1 were prepared from E. coli and human HeLa cells, respectively. Direct binding between the two recombinant proteins, following dephosphorylation of CLK1 by PP1, was assessed in vitro using a GST pulldown assay. The RIOK2-CLK1 interaction was not significantly affected by the pretreatment of CLK1 with PP1 (Figures S5G-S5I). These results suggest that PP1-mediated dephosphorylation of CLK1 does not significantly affect RIOK2 binding. To examine the second possibility, RIOK2 immobilized on beads was incubated with CLK1 at either 37□ or 42□, and the complexes were subsequently recovered. Direct binding of RIOK2 to CLK1 was enhanced at 42□ compared to at 37□ (Figures 5C and 5D), whereas the binding of CLK1 to its substrate SRSF9 remained unaffected by temperatures (Figures S5J-S5L). These data indicate that RIOK2-CLK1 interaction is thermo-dependent, showing enhanced association under thermal stress. In cells, the interaction weakens during stress recovery (Figures 5A and 5B). Intriguingly, this reduction is partially prevented by RIOK2 inhibition (Figures 5E and 5F), suggesting that their interactions during stress recovery are regulated not only by temperature changes but also RIOK2-dependent phosphorylation of CLK1. During recovery, CLK1 remains tightly associated with RIOK2 until it is phosphorylated, which may also contribute to the rapid rephosphorylation of CLK1.

We also assessed whether RIOK2 kinase activity is temperature-dependent. Using recombinant RIOK2 and CLK1 proteins, we found that neither RIOK2 autophosphorylation nor its ability to phosphorylate CLK1 was significantly affected at 42□ compared to 37□ in vitro (Figures 5G and 5H). To examine RIOK2 activity under thermal stress in cells, we exposed cells to 42□ for 1 hour, which induced CLK1 dephosphorylation. While maintaining thermal stress, cells were treated with calyculin A alone or together with the RIOK2 inhibitor, CQ211 (Figure 5I). Calyculin A treatment restored CLK1 phosphorylation, indicating that some protein kinases remain active under thermal stress (Figure. 5J). In contrast, co-treatment with calyculin A and CQ211 abolished this phosphorylation, confirming that it is dependent on RIOK2 activity (Figure. 5J). These results suggest that RIOK2 retains its kinase activity under thermal stress. Notably, although RIOK2 remains active, CLK1 is dephosphorylated under these conditions, suggesting that PP1 is activated during thermal stress and counteracts RIOK2 in regulating CLK1 phosphorylation, even though RIOK2 binds more efficiently to CLK1 under these conditions. Taken together, these findings indicate that under thermal stress, RIOK2 remains catalytically active and interacts more tightly with CLK1, which may prime CLK1 for the rapid rephosphorylation during stress recovery (Figure 5K).

### PPP1R2 acts as a reversible thermosensor controlling PP1 activity to regulate CLK1 phosphorylation

As described above, the interaction between PP1 and CLK1 was significantly enhanced during thermal stress compared to normal and recovery conditions (Figures 5A and 5B). Since PP1 specifically dephosphorylates CLK1 during thermal stress, this increased interaction is consistent with its functional role. However, given that PP1 also binds to CLK1 under normal and recovery conditions (Figures 5A and 5B), yet CLK1 is markedly dephosphorylated only during thermal stress (Figures 3A and 3B), these observations suggest that PP1 is specifically activated in response to thermal stress. PP1 activity is regulated by a lerge set of regulatory subunits, with more than 200 identified to date.^46^ Based on this, we hypothesized that the interaction between PP1 and specific regulatory subunits may change in response to temperature shifts. To test this, we evaluated the interaction between PP1 and several nuclear regulatory subunits under different temperature conditions using PPP1CC coIP-WB, focusing on PPP1R2, PPP1R8, and PPP1R10. We found that PPP1R2 dissociates from PP1 significantly under thermal stress and reassociates duringl stress recovery (Figures 6A, B). In contrast, the interactions between PP1 and PPP1R8 or PPP1R10 were largely unaffected by temperature changes. As PPP1R2 is one of the earliest characterized nuclear inhibitors of PP1, these findings suggest that it functions as a thermosensitive inhibitor, with its dissociation under thermal stress leading to PP1 activation. Interestingly, PPP1R2 has been reported as a heat-stable protein with an entirely intrinsically disordered structure (Figure S6A), ^47,48^ supporting its potential role in thermosensing.

**Figure 6.**
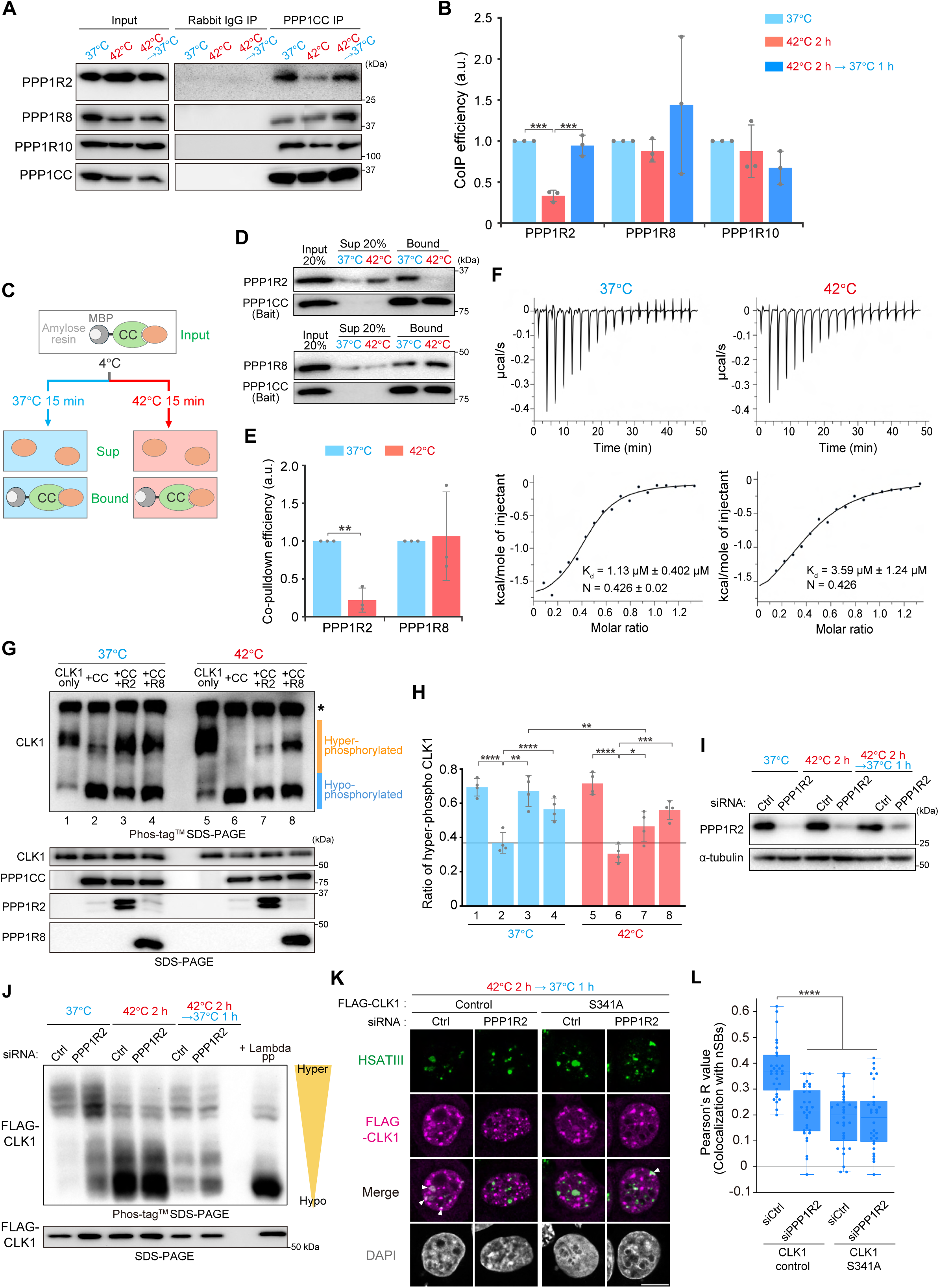
PPP1R2 functions as a reversible thermosensor regulating PP1 activity and CLK1 phosphorylation status. (A) Interactions of PPP1CC with PPP1R2/PPP1R8/PPP1R10 across temperature conditions assessed by coIP-WB. (B) Quantification of coIP efficiency of PPP1R2/PPP1R8/PPP1R10 in (A). CoIP efficiency was calculated as the ratio of PPP1CC IP band intensity of PPP1R2/PPP1R8/PPP1R10 to that of PPP1CC. Data are presented as mean ± SD (n = 3). ***0.0001 < *p* < 0.001 (Tukey’s multiple comparisons test). (C) Schematic of the experimental design to assess temperature-dependency of PPP1CC interaction with PPP1R2 or PPP1R8. (D) Direct interactions of recombinant PPP1CC with PPP1R2/PPP1R8 at 37□ and 42□□ assessed by MBP pulldown and WB. (E) Quantification of co-pulldown efficiency of PPP1R2/PPP1R8 in (D). Co-pulldown efficiency was calculated as the ratio of the band intensity of PPP1R2/PPP1R8 to that of CLK1 in the bound fraction. Data are presented as mean ± SD (n = 3). **0.001 < *p* <0.01 (two-tailed t-test with Bonferroni correction). (F) ITC results were obtained by titrating of PPP1R2 into the solution of PPP1CC at 37□ and 42□. Dissociation constants (Kd) and stoichiometry values (N) are shown in the lower panel. (G) Phos-tag™ SDS-PAGE and WB analysis of CLK1 after incubation with recombinant PPP1CC alone or with PPP1R2/PPP1R8 (upper panel). Hyper- and Hypo-phosphorylated forms are indicated at the right. Asterisks denote putative aggregated CLK1 proteins. Recombinant proteins were verified by standard SDS-PAGE and WB (lower panel). (H) Quantification of the proportion of hyper-phosphorylated CLK1 in (G). Each bar corresponds to the lane numbers shown in (G). Data are presented as mean ± SD (n = 4). \*\*\*\**p* < 0.0001, ***0.0001 < *p* < 0.001, **0.001 < *p* <0.01, *0.01 < *p* < 0.05 (Šidák’s multiple comparisons test). (I) Western blot validation of PPP1R2 knockdown efficiency. α-tubulin is a loading control. (J) Effect of PPP1R2 knockdown on FLAG-CLK1 phosphorylation status, assessed by Phos-tag™ SDS-PAGE (upper panel) and standard SDS-PAGE (lower panel) followed by WB. siRNAs (20 nM) are indicated above the upper panel. (K) Subnuclear localization of CLK1 during stress recovery visualized by HSATIII RNA-FISH combined with FLAG IF. siRNAs (20 nM) are indicated above the panel. Scale bar, 10 μm. (L) Quantification of CLK1 colocalization with nSBs (n = 30) in (K). \*\*\*\**p* < 0.0001 (Dunn’s multiple comparisons test).

To assess whether the PPP1CC-PPP1R2 interaction is thermosensitive, we performed an in vitro binding assay using purified recombinant proteins (Figure S6B). We confirmed that PPP1R2, PPP1R8 (used as a control), and CLK1 directly bind to PPP1CC (Figures S6C-S6E). To examine the temperature dependence of their interaction, PPP1CC was incubated with either PPP1R2 or PPP1R8, and the resulting complexes were purified at 4□ using MBP-tag on PPP1CC. The purified complexes were subsequently incubated at 37□ or 42□ (Figure 6C). PPP1CC-PPP1R2 complexes remained stable at 37□ but clearly dissociated at 42□. In contrast, PPP1CC-PPP1R8 remained stable at both temperatures (Figures 6D and 6E). These results are consistent with the observation that PPP1R2 dissociates from PPP1CC upon thermal stress in cells (Figure 6A), indicating that the PPP1CC-PPP1R2 interaction is thermosensitive. To quantitatively assess this thermosensitivity, we determined the dissociation constant (Kd) of the PPP1CC-PPP1R2 interaction using isothermal titration calorimetry (ITC). The Kd at 42□ (3.59 μM) was approximately three times higher than that at 37□ (1.13 μM) (Figure 6F). Based on the Kd values, the occupancy of the PPP1CC-PPP1R2 complex can be estimated under physiological concentrations. According to the OpenCell database,^49^ cellular concentrations of PPP1CC and PPP1R2 are 380 nM and 50 nM, respectively. At 37□, with a Kd of 1.13 μM, approximately 25% of PPP1R2 is bound to PPP1CC. In contrast, at 42□, where the Kd increases to 3.59 μM, the bound fraction drops to 11%. Thus, the complex is approximately 2.3-fold less likely to form at 42□ compared with 37□, indicating that their interaction is highly sensitive to elevated temperature.

To verify that PPP1R2 dissociation from PP1 leads to PP1 activation, we performed an in vitro phosphatase assay using phosphorylated recombinant CLK1. First, we confirmed that phosphorylated CLK1 is dephosphorylated by PPP1CC in a dose-dependent manner (Figure S6F). Moreover, both PPP1R2 and PPP1R8 inhibited PPP1CC-mediated dephosphorylation of CLK1, confirming their roles as inhibitory subunits of PP1 in this system (Figures S6G and S6H). We then evaluated whether the inhibitory activities of PPP1R2 and PPP1R8 differ between 37□ and 42□.

PPP1CC efficiently dephosphorylated CLK1 at both temperatures in the absence of inhibitory subunits (Figure 6G, lanes 1, 2, 5 and 6; Figure 6H). At 37□, dephosphorylation was inhibited in the presence of either PPP1R2 or PPP1R8 (Figure 6G, lanes 3 and 4; Figure 6H). In contrast, at 42□, CLK1 was significantly dephosphorylated even in the presence of PPP1R2 (Figure 6G, lane 7; Figure 6H), suggesting that PP1 is activated by dissociation of PPP1R2 at elevated temperature. PPP1R8, however, continued to inhibit CLK1 dephosphorylation at 42□ (Figure 6G, lane 8; Figure 6H). Next, we examined the role of PPP1R2 in cells by evaluating CLK1 phosphorylation and its localization to nSBs upon PPP1R2 knockdown (Figure 6I). PPP1R2 knockdown promoted CLK1 dephosphorylation under both normal and recovery conditions (37□ and 42□→37□) (Figure 6J) and significantly prevented CLK1 localization to nSBs during stress recovery (Figures 6K and 6L). Importantly, PPP1R2 knockdown had no additional effect on the CLK1 S341A mutant (Figures 6K and 6L). These results are consistent with the in vitro evidence that the thermosensitive PPP1CC-PPP1R2 interaction regulates CLK1 phosphorylation at Ser341, which is required for its nSB localization. Thus, we conclude that PPP1R2 functions as a reversible thermosensor that transiently activates PP1 during thermal stress, thereby controlling the temperature-dependent phosphorylation and subnuclear localization of CLK1.

### CLK1 phosphorylation status is dynamically regulated through temperature-dependent changes in the balance between PP1 and RIOK2 activities

To dissect the temperature-dependent regulation of CLK1 phosphorylation by RIOK2 and PP1, we reconstituted phosphorylation status of CLK1 in vitro using purified recombinant proteins. We first investigated whether RIOK2, PPP1CC, and CLK1 form a trimeric complex. Recombinant proteins were mixed and sequentially purified: first by GST-tagged RIOK2 pulldown, followedby MBP-tagged PPP1CC pulldown. If a stable trimeric complex formed, CLK1 would be detected in the final eluate (Figure 7A, Bound-2). However, CLK1 was barely detectable after the second purification, suggesting that a stable trimeric complex does not form (Figure 7B).

**Figure 7.**
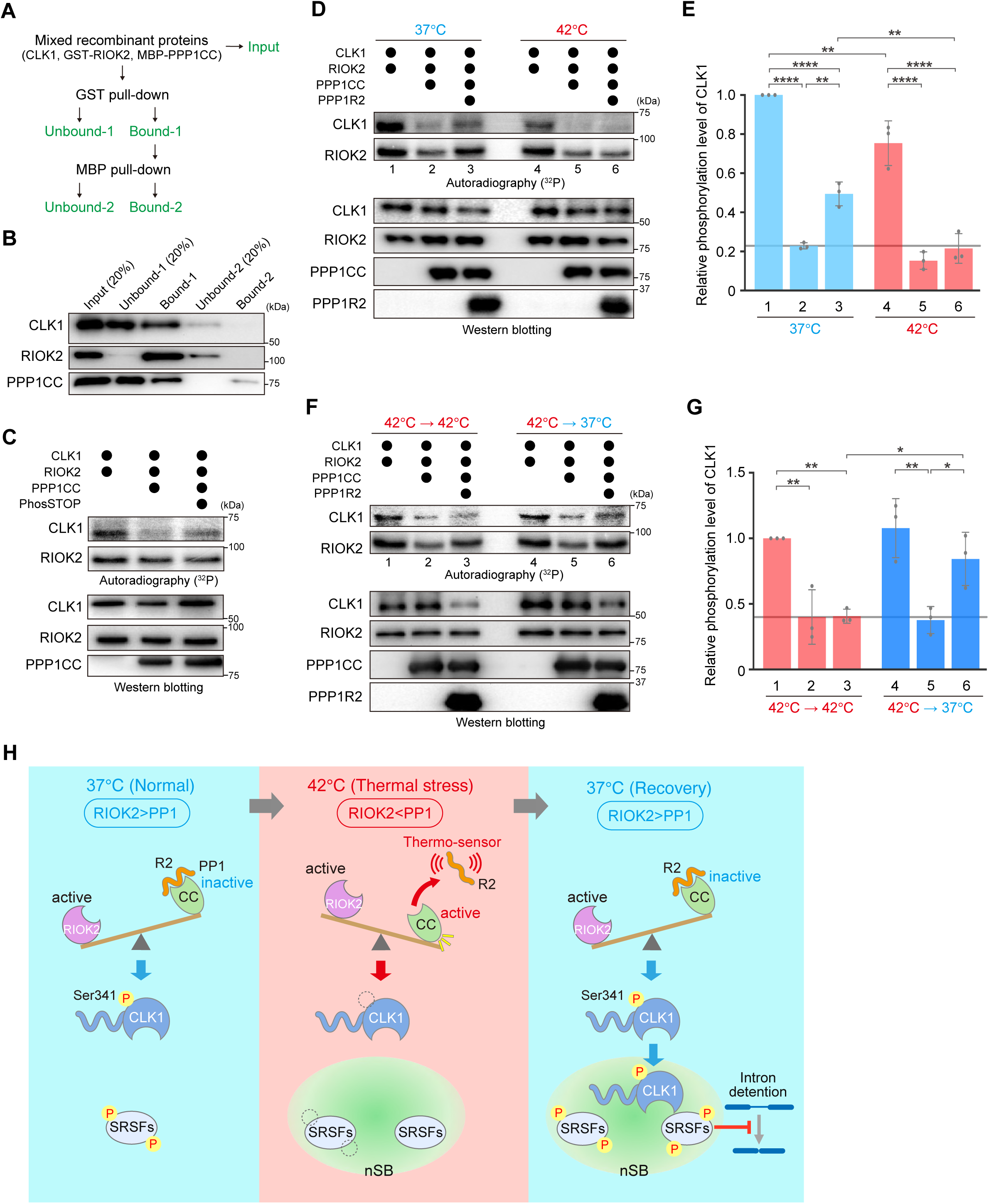
Temperature-dependent balance between PP1 and RIOK2 activities determines CLK1 phosphorylation status. (A) Schematic of the experimental workflow to test whether RIOK2, CLK1, and PPP1CC form a ternary complex. (B) WB of the two-step pulldown experiment shown in (A). (C) In vitro phosphorylation of RIOK2 and CLK1 with and without PPP1CC assessed by autoradiography with WB validation. PhosSTOP^TM^ is used as an inhibitor of PPP1CC. (D, F) In vitro phosphorylation of RIOK2 and CLK1 with PPP1CC alone or combined with PPP1R2 (D) at 37□ and 42□, or (F) at 42□→42□ or 42□→37□ analyzed by autoradiography with WB validation. (E, G) Quantification of CLK1 autoradiographic signals in (D, F). Each bar corresponds to the lane numbers shown in (D, F). Data are presented as mean ± SD (n = 3). \*\*\*\**p* < 0.0001, **0.001 < *p* <0.01, *0.05 < *p* <0.01 (Šidák’s multiple comparisons test). (H) Model for the molecular mechanism of temperature-dependent CLK1 localization to nSBs through thermosensing CLK1 phosphorylation.

Next, we investigated how the coordinated actions of RIOK2 and PP1 regulate CLK1 phosphorylation in a temperature-dependent manner. Using autoradiography with [γ-^32^P] ATP, we monitored CLK1 phosphorylation under different temperature conditions. In this system, Ser341 of CLK1 was directly phosphorylated by RIOK2 (Figures 3H, 3I, S3E, and S3F). The addition of PPP1CC reduced CLK1 phosphorylation, which was partially rescued by the phosphatase inhibitor (PhosSTOP^TM^) (Figure 7C), indicating that PP1 antagonizes RIOK2-mediated phosphorylation in a phosphatase activity-dependent manner. We then examined the combined effect of RIOK2, PPP1CC, and PPP1R2 on CLK1 phosphorylation and its temperature dependency. RIOK2 efficiently phosphorylated CLK1 at both temperatures, and this phosphorylation was suppressed by PPP1CC (Figure 7D, lanes 1, 2, 4, and 5; Figure 7E). Notably, in the presence of PPP1R2, CLK1 phosphorylation by RIOK2 became thermosensitive: it was detectable at 37□ but markedly reduced at 42□, despite RIOK2 retaining its kinase activity at the higher temperature (Figure 7D, lanes 3 and 6; Figure 7E). To mimic the temperature changes during stress recovery, when CLK1 is phosphorylated and localizes to nSBs, the recombinant proteins were first incubated at 42□ and then either maintained at 42□ or shifted to 37□. CLK1 phosphorylation was higher under the recovery condition (42□→37□) than under continuous thermal stress (42□→42□), but only in the presence of PPP1R2 (Figures 7F and 7G). These results indicate that our in vitro system successfully recapitulates the thermosensitive CLK1 phosphorylation observed in cells during the thermal stress response. Moreover, RIOK2, PPP1CC, and PPP1R2 are sufficient to reconstitute this regulation, at least in part. Taken together, CLK1 phosphorylation, and thus its subcellular localization, is dynamically regulated by temperature-dependent changes in the balanced interplay between RIOK2 kinase activity and PPP1R2-modulated PP1 phosphatase activity (Figure 7H).

## DISCUSSION

nSBs control temperature-dependent splicing of >400 pre-mRNAs in human cells. For a subset, they act as a reaction crucible for SRSF phosphorylation: SRSFs accumulate in nSBs during thermal stress, and subsequently, their kinase CLK1 is recruited during recovery. This spatiotemporal coordination enables rapid SRSF phosphorylation that promotes intron detention in target pre-mRNAs. A key regulatory step is temperature-dependent CLK1 localization to nSBs, however the molecular mechanism remained unclear. Here, we show that CLK1 Ser341 phosphorylation is thermo-regulated and essential for nSB targeting: dephosphorylated during stress and rapidly rephosphorylated during recovery. Mass spectrometry and mutational analysis identified Ser341 as critical for nSB targeting and intron detention by nSBs (Figure 2K). Ser341 is conserved across CLK family but substituted only in CLK3, which is not under RIOK2 regulation (Figures S3J-S3L), suggesting division of labor among CLK family members. However, the mechanism by which phosphorylated CLK1 specifically localizes to nSBs remains unknown, and identifying the responsible factor(s) will be an important future direction. Together, our findings further expand the understanding of the multilayered mechanisms underlying the thermoregulation of CLK1, encompassing its gene expression,^3^ enzymatic activity,^50^ and, as demonstrated here, subcellular localization.^22^

The nSB-LRs within the N-terminal IDR of CLK1 are essential for its its localization to nSBs during recovery. Interactome analysis of the nSB-LRs identified two counteracting enzymes that regulate CLK1 Ser341 phosphorylation: RIOK2 phosphorylates CLK1, and PP1 dephosphorylates it during thermal stress. Interestingly, RIOK2 does not affect CLK1 localization to nuclear speckles during recovery (Figures S2P-S2R), indicating compartment-specific control of RIOK2, although the precise subnuclear site where RIOK2 phosphorylates CLK1 remains unresolved. PP1 dephosphorylates CLK1 Ser341 and prevents its localization to nSBs. PP1 also dephosphorylates SRSFs during stress,^33^ suggesting that a common enzyme regulates dephosphorylation of both substrates (SRSFs) and the enzyme (CLK1) to ensure the crucible function of nSBs. However, their localization outcomes differ: dephosphorylated SRSFs, but not dephosphorylated CLK1, can enter nSBs, enabling their sequential localization in response to temperature shifts. Notably, RIOK2 inhibition and PP1 overexpression do not fully inhibit CLK1 localization to nSBs, leaving open the possibility of additional regulatory pathways.

Phosphorylation status of CLK1 Ser341 is determined by temperature-dependent changes in the balance between RIOK2 and PP1 activities, with PPP1R2 as a key regulatory factor (Figure 7H). At 37°C, PPP1CC is inhibited through direct association with PPP1R2, whereas under thermal stress (42°C), PPP1R2 dissociates, activating PP1. In vitro binding and ITC analysis revealed a threefold increase in Kd under stress, indicating their thermosensitive interaction. Previous study proposed that PPP1R8 acts as a thermosensitive PP1 inhibitor, as it dissociates from PP1 under thermal stress and its overexpression prevents PP1-mediated SRSF10 dephosphorylation.^33^ In contrast, our analyses show that PPP1R8 barely dissociate from PP1 under thermal stress, and the PPP1CC-PPP1R8 interaction is not thermosensitive in vitro (Fig. 6A–6E). Instead, our data strongly indicate that PPP1R2 functions as an essential thermosensor for CLK1 dephosphorylation specifically under thermal stress. During recovery, PPP1R2 reassociates with and inactivates PP1, promoting CLK1 phosphorylation, its localization to nSBs, and nSB-mediated splicing regulation. These results establish PPP1R2 as a key regulator acting as a thermosensor controlling the nSB crucible mechanism. Although, comprehensive identification of thermosensitive PP1 regulatory subunits has not been performed, additional factors may also function as thermosensors. PPP1R2, an intrinsically disordered protein without folded domains, resembles heat-resistant obscure (Hero) proteins. Some of which function in a chaperone-like manner to solubilize protein aggregation in cells.^48^ Our results represent a novel functional class of highly disordered proteins. Moreover, understanding the detailed molecular mechanism of PPP1R2’s thermosensing function will provide broader insight into the roles of highly disordered proteins.

While PP1 activity changes with temperature, RIOK2 kinase activity remains constant between 37°C and 42°C (Figures 5G and 5H). This suggests that CLK1 Ser341 phosphorylation, its nSB localization, and splicing regulation by nSBs are determined by temperature-dependent changes in the balance of activities between RIOK2 and PP1, primarily controlled by PPP1R2 (Figure 7H). During thermal stress, PP1 dominates, thereby dephosphorylating CLK1 and preventing its nSB localization. During recovery, PP1 is inhibited by PPP1R2, allowing RIOK2 to phosphorylate CLK1. Our in vitro system indicates that these four recombinant proteins are sufficient to recapitulate thermosensitive CLK1 phosphorylation (Figures 7D-7G). Notably, the RIOK2-CLK1 interaction is thermo-regulated, strengthening under thermal stress, as shown by our coIP-WB and in vitro binding assay (Figures 5A-5D). Since RIOK2 phosphorylates CLK1 during recovery, this strengthening was unexpected and suggests that the RIOK2 association alone is insufficient for CLK1 phosphorylation in the presence of active PP1. However, additional regulatory factors may also modulate RIOK2 activity during temperature shifts. The enhanced interaction likely serves as a priming mechanism, enabling CLK1 rapid phosphorylation once temperature returns to normal and PP1 is inactivated by PPP1R2. Together, these findings reveal a multilayered thermosensing mechanism that sharply regulates temperature-dependent pre-mRNA splicing via the nSB crucible function.

### Limitations of the study

PPP1R2 was identified as a thermosensor for PP1-mediated dephosphorylation. However, evaluating whether its dissociation drives global protein dephosphorylation is technically challenging, as PPP1R2 itself dissociates under thermal stress, making simple depletion inappropriate; a PPP1R2 mutant that remains bound to PP1 even at 42□ would be required. Moreover, the intrinsically disordered nature of PPP1R2 complicates mechanistic analysis underlying this process. Demonstrating the functional significance of RIOK2 preloading onto CLK1 is also difficult, since selectively inhibiting their interaction during thermal stress without affecting it during recovery is technically challenging. How phosphorylated CLK1 is recruited to nSBs, and whether this step itself is thermosensitive, remains unclear. The mechanism may be elucidated by further analyses of nSB-localized proteins that specifically interact with CLK1 phophorylated at Ser341. Finally, the physiological roles of nSB-mediated splicing, particularly, the significance of this primate-specific membraneless organelle in stress responses and its potential role in protecting fragile tissues, remain to be clarified.

## RESOURCE AVAILABILITY

### Lead contact

Further information and requests for resources and reagents should be directed to and will be fulfilled by the Lead Contact, Tetsuro Hirose.

### Materials availability

All unique plasmids and constructs generated in this study are available from the Lead Contact.

### Data and code availability

All data reported in this paper will be shared by the Lead Contact upon reasonable request. The mass spectrometry raw data were deposited in the Japan ProteOme STandard Repository (jPOST) under the ID JPST004102. No original code was generated.

## ACKNOWLEDGMENTS

We thank the members of the Hirose laboratory for their valuable discussions and support.We also thank FBS Core Facility at The University of Osaka for technical assistance. This research was supported by grants from JST CREST (grant no. JPMJCR20E6 to T.H., JPMJCR23B3 and JPMJCR20E6 to S.A., JPMJCR20E3 to N.N.N.), AMED (grant no. 21479280 to T.H., and CREST grant no. 23gm1610010h0002 to S.A.), JSPS KAKENHI (grant no. 21H05276 and 24K21933 to T.H., 22H02225 to S.A., 25H01321 to N.N.N.), and National Cancer Center Research and Development Funds (no. 2023-A-01 to S.A.). T.U. was supported by JST SPRING and JSPS Fellowship DC2 (25KJ1741).

## AUTHOR CONTRIBUTIONS

T.U., K.N., and T.H. conceived and designed this study. T.U. conducted most of the experiments.

S.A. performed the mass spectrometry analyses, and N.N.N. carried out the ITC measurements. I.T. provided technical support for the in vitro experiments. T.U., K.N., and T.H. wrote the manuscript. All authors have reviewed the manuscript.

## SUPPLEMENTAL INFORMATION

### Document S1. Figures S1-S6 and Supplemental References (PDF)

**Table S1. Interactions between CLK1 and candidate regulators or catalytic subunits of PP1 detected by mass spectrometry, related to Figures 2 and 4 (Excel file).**

Interaction score between 7 candidate regulators or catalytic subunits of PP1 and CLK1 deletion mutants (Figure 1A and S1E). Data are shown as relative amounts of proteins immunoprecipitated with vector control or CLK1 deletion mutants of two biological replicates, with each sample measured twice by mass spectrometry and averaged.

**Table S2. The list of primers used in this paper, related to STAR Methods (Excel file).** Oligonucleotide primers used for cloning, mutagenesis, and qRT-PCR in this study. All sequences are written in the 5’→3’ orientation.

## STAR★METHODS

### KEY RESOURCES TABLE

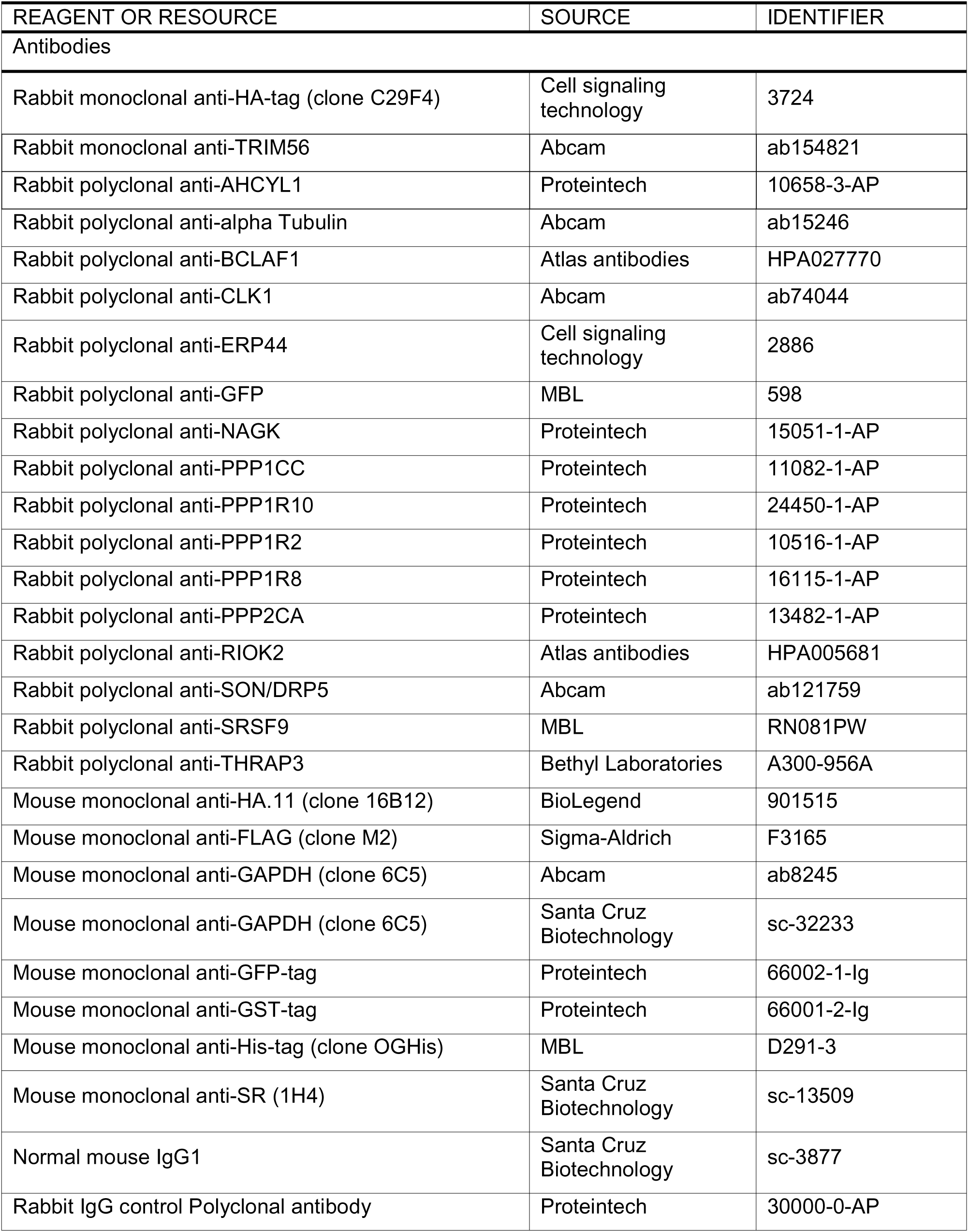
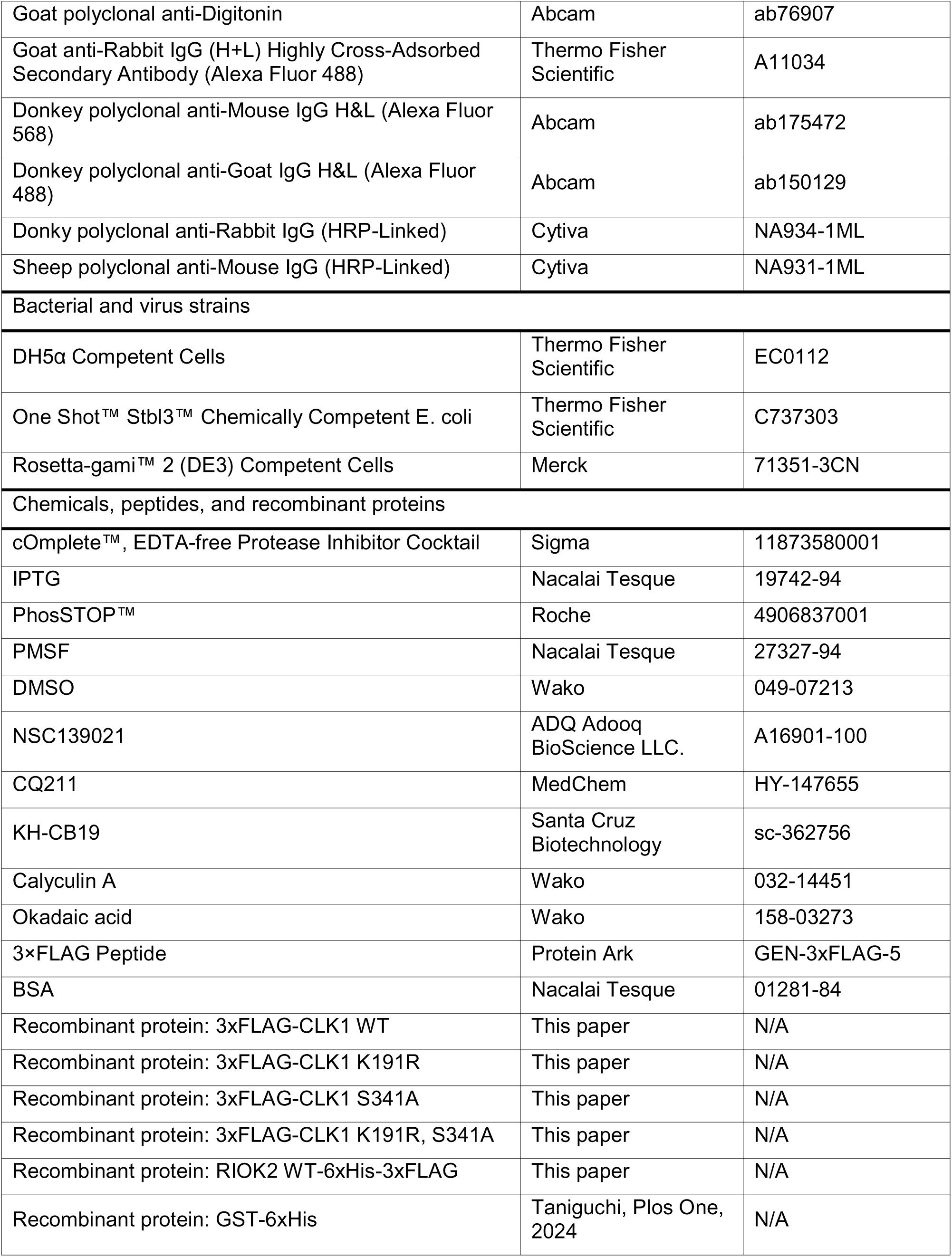
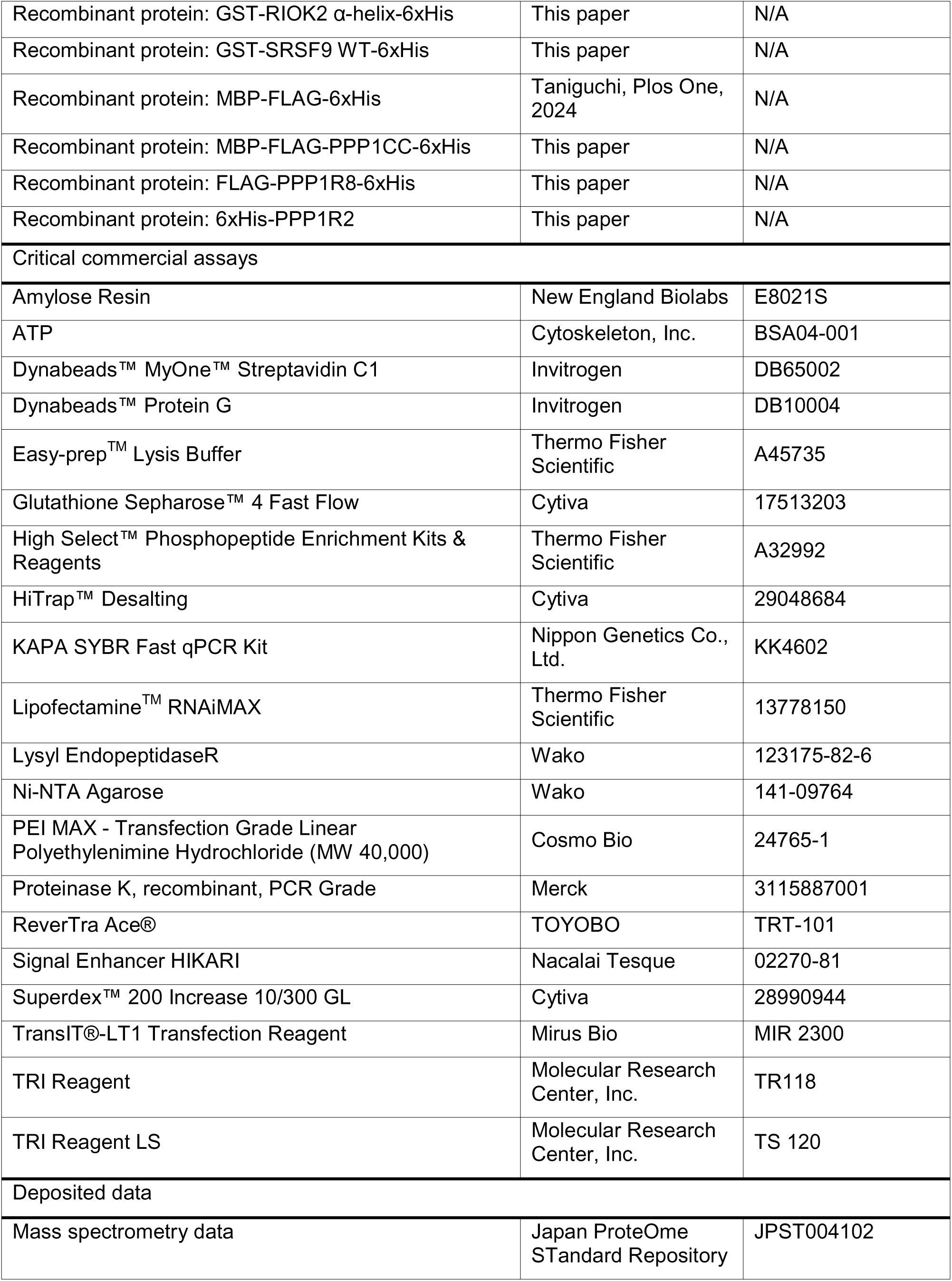
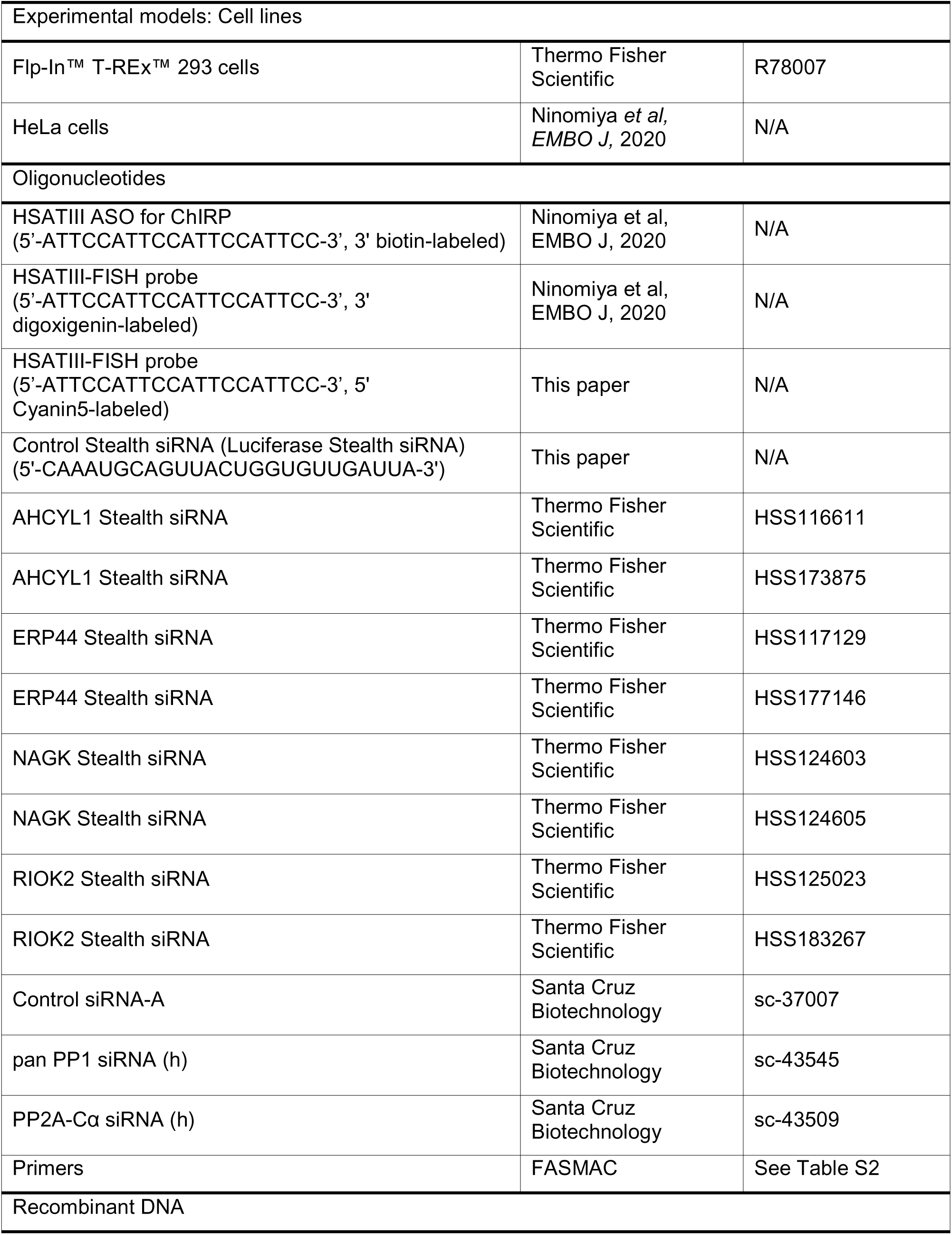
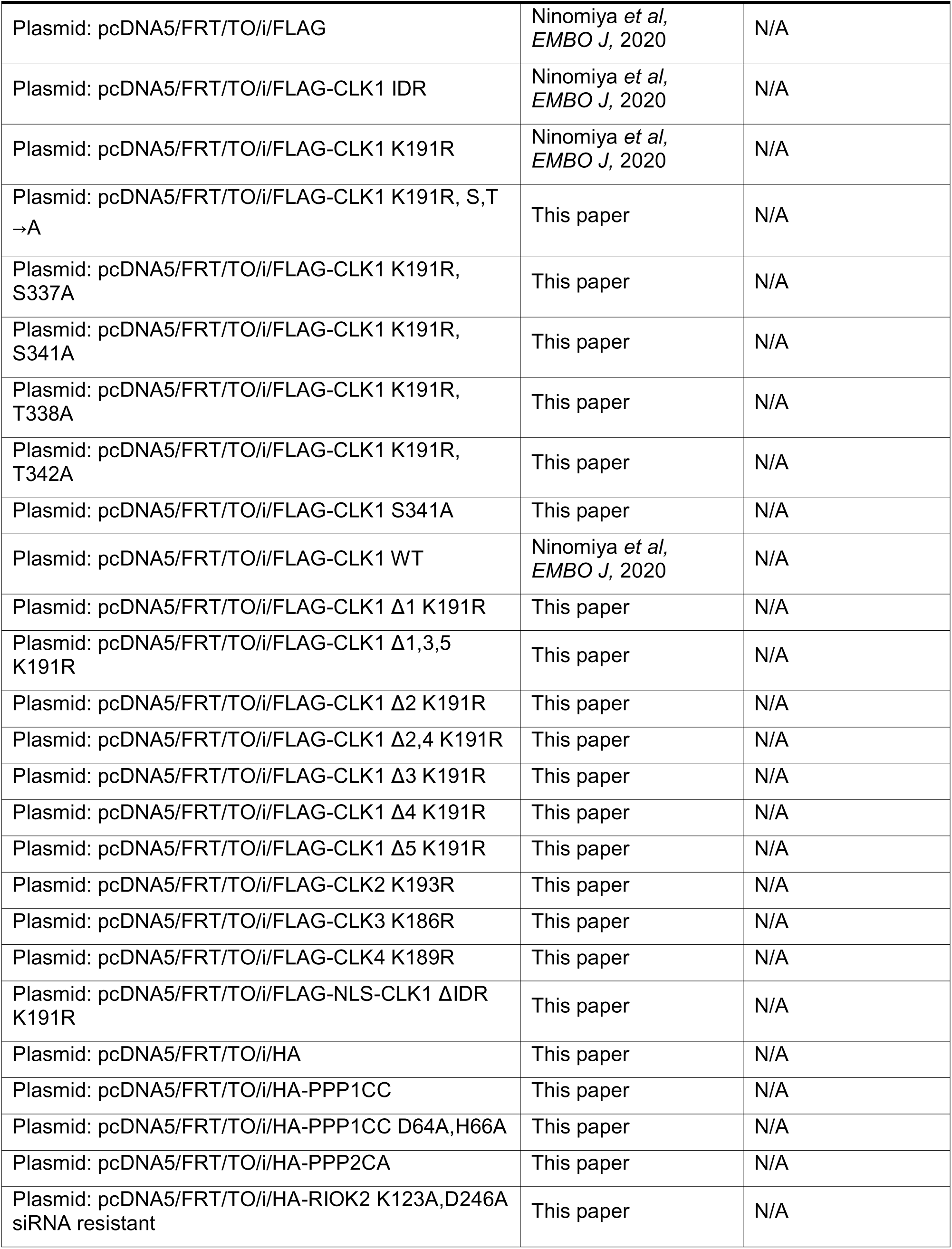
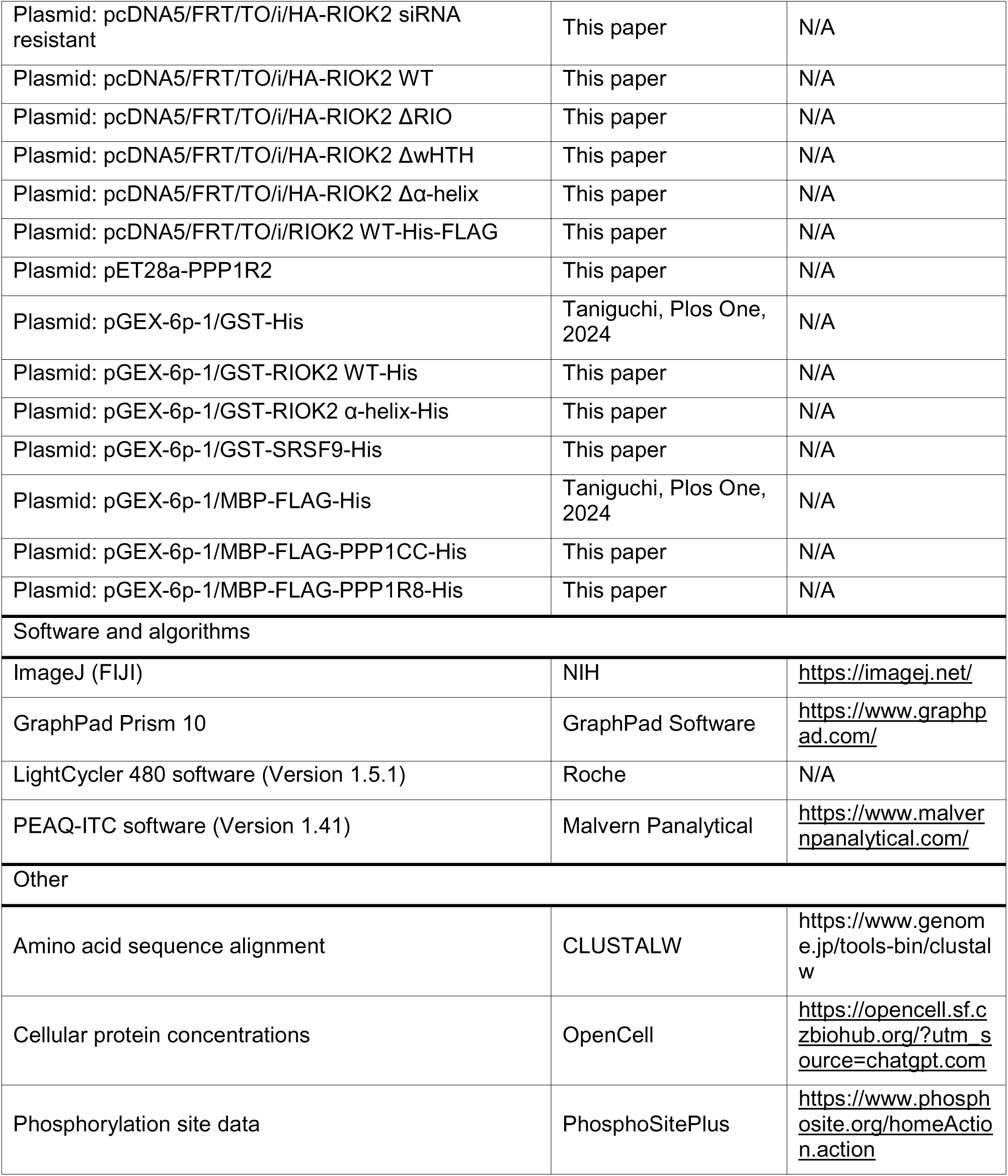

#### METHOD DETAILS

##### Cell culture and chemical treatment

HeLa cells were cultured in DMEM (Nacalai Tesque) with 10% fetal calf serum (Sigma) and antibiotics (100 U/ml streptomycin, 100 μg/ml penicillin; Nacalai Tesque) at 37□, 5% CO_2_. For thermal stress, cells were incubated at 42□ in 5% CO_2_. For chemical treatment, medium was replaced with fresh pre-warmed medium containing chemical compounds. Used chemical compounds are listed in Key resources table.

### Plasmid construction

DNA fragments were PCR-amplified from Human cDNAs and inserted into the tag-inserted pcDNA5/FRT/TO/i vectors^22^ for expression in cultured cells. For E. coli expression, the fragments were inserted into pET28a or pGEX-6p-1.^51^ Deletion, amino acid substitution, and siRNA-resistant mutations were introduced via inverted PCR or In-Fusion® Snap Assembly (TaKaRa). Primers are listed in Table S2.

### Plasmid transfection and knockdown using siRNA

Plasmids were transfected as described (Ninomiya et al, 2020, 2021). For siRNA knockdown, Stealth RNAi™ siRNAs (Thermo Fisher Scientific) were transfected at 20 nM or 10 nM each (for double knockdown), using LipofectamineTM RNAiMAX (Thermo Fisher Scientific) 48 h before assays. After 6 h, the medium was replaced, and plasmids were transfected 24 h before the assay. Plasmids and siRNAs are listed in Key resources table.

### RNA-FISH and immunofluorescence

RNA-FISH and immunofluorescence were performed as reported (Ninomiya et al, 2020, 2021), with minor modifications. For HSATIII detection, Dig-labeled or Cy5-labeled HSATIII antisense oligo was used. Images were acquired using an LSM900 confocal microscope (Zeiss) and analyzed using ImageJ (FIJI). Used antibodies are listed in Key resources table.

### Western blotting

Western blotting and Phos-tag™ SDS-PAGE were performed as described (Ninomiya et al, 2020, 2021). If necessary, signals were enhanced using Signal Enhancer HIKARI (Nacalai Tesque). Images were detected using ChemiDoc Touch (Bio-Rad) and analyzed with ImageJ (FIJI). For Lambda protein phosphatase treatment, cells expressing FLAG-tagged CLK1 were washed with PBS and lysed in NEBuffer for PMP (NEB) containing 0.1% NP-40. After sonication, samples were centrifuged at 13,000 g for 1 min at 4□, and Lambda protein phosphatase (NEB) and 1 mM MnCl_2_ were added. The samples were incubated at 30□ for 1 h. Used antibodies are listed in Key resources table.

### ChIRP

HSATIII-ChIRP was performed as described previously (Ninomiya et al, 2020, 2021). Briefly, cells were crosslinked with 1% PFA/PBS for 30 min, quenched with 150 mM glycine, washed with ice-cold PBS, and collected. The collected cells were lysed and sonicated. Insoluble fraction was removed, and lysates were then hybridized with biotinylated HSATIII antisense oligo and captured with Dynabeads MyOne Streptavidin C1 (Thermo Fisher Scientific). Beads were washed, and bound proteins were eluted with SDS sample buffer and analyzed by SDS–PAGE and western blotting. Bound RNAs were de-crosslinked by proteinase K treatment (Merck) and heating, then purified using TRI Reagent LS (Molecular Research Center, Inc.).

### Immunoprecipitation

Cells were washed with ice-cold PBS twice and lysed in IP-150 buffer (50 mM Tris-HCl pH 7.5, 150 mM NaCl, 0.5% NP-40) supplemented with cOmplete™ Protease Inhibitor Cocktail (Sigma), PhosSTOP™ (Roche), and 1 mM PMSF. After sonication, the lysates were centrifuged at 13,000 g for 1 min at 4□ or at 17,800 g for 10 min at 4□. The supernatant was incubated with antibody-conjugated Dynabeads™ Protein G (Invitrogen) and rotated at 4□ for 1-2 h. Beads were washed five times with IP-150 buffer, and proteins were eluted using 5x SDS sample buffer or 3x FLAG Peptide (Protein Ark).

### Mass Spectrometry-Based Preparation, Measurement, and Data Analysis

For immunoprecipitation (IP) samples, proteins were precipitated with trichloroacetic acid (TCA), washed with chilled acetone, and dissolved in 7 M guanidine hydrochloride. The solubilized proteins were reduced with tris(2-carboxyethyl)phosphine (TCEP), alkylated with iodoacetamide (IAA) to carbamidomethylate cysteine residues, and digested sequentially with lysyl endopeptidase (Wako) and trypsin (Thermo Fisher Scientific). For phosphoproteome analysis, cells were lysed in Easy-prep™ Lysis Buffer (Thermo Fisher Scientific) supplemented with PhosSTOP™ (Roche), followed by protein precipitation with acetone. The resulting pellets were dissolved in guanidine hydrochloride and processed in the same way as the IP samples: reduced with TCEP, alkylated with IAA, and digested with lysyl endopeptidase and trypsin. Phosphopeptides were enriched using the High-Select™ Fe-NTA Phosphopeptide Enrichment Kit (Thermo Fisher Scientific) according to the manufacturer’s instructions. All resulting peptides were analyzed using an Evosep One LC system (EVOSEP) coupled to a Q-Exactive HF-X mass spectrometer (Thermo Fisher Scientific). Chromatography was performed with 0.1% formic acid in water (solution A) and 0.1% formic acid in 99.9% acetonitrile (solution B) as mobile phases. Data were acquired in data-dependent acquisition mode, selecting the top 25 precursor ions within the m/z range of 380–1500. MS/MS spectra were searched against the human Swiss-Prot protein databases using Proteome Discoverer 2.5 with the SEQUEST search engine, and filtered at a 1% false discovery rate (FDR) for peptide-spectrum matches. The data have been deposited to the jPOST repository under accession number JPST004102.

### RNA extraction and quantitative RT-PCR

Total RNAs were extracted from cultured cells or ChIRP samples, using TRI Reagent or TRI Reagent LS (Molecular Research Center, Inc.) according to the manufacturer’s protocol. RNA was treated with RQ1 DNase (Promega) and reverse-transcribed using ReverTra Ace® (TOYOBO). The sample and reference cDNAs were amplified using the KAPA SYBR Fast qPCR Kit (Nippon Genetics Co., Ltd.) and quantified with the LightCycler 480 SystemII (Roche). Primers are listed in Table S2.

### Expression and purification of recombinant proteins

For recombinant protein expression in E. coli, Rosetta-gami™ 2 (DE3) competent cells (sigma) were transformed with plasmids, and expression was induced with 0.5 mM IPTG at OD600 = 0.6, followed by incubation at 30□ for 4 h or 20□ overnight. Cells were pelleted and lysed in His-tag purification buffer (20 mM Tris-HCl pH 7.5, 500 mM NaCl, 20 mM imidazole, 10% glycerol) supplemented with 1% NP-40 and 1 mM PMSF. After lysozyme treatment (0.2 mg/ml, Wako), lysates were sonicated and centrifuged. His-tagged proteins were purified using Ni-NTA Agarose (Wako) and eluted with imidazole. If necessary, GST- or MBP-tagged proteins were further purified using Glutathione Sepharose™ 4 Fast Flow (Cytiva) or Amylose Resin (NEB), respectively. Finally, proteins were dialyzed into dialysis buffer (20 mM Tris-HCl pH 7.5, 500 mM NaCl, 1 mM DTT, 0.2 mM EDTA, 10% glycerol).

For recombinant protein expression in Flp-In™ T-REx™ 293 cells, expression was induced with 1 μg/ml doxycycline, while HeLa cells were transfected with PEI Max™ (Cosmo Bio Co., Ltd.). For purification, cells were pelleted and lysed in IP-1000 buffer (20 mM Tris-HCl pH 7.5, 1 M NaCl, 0.5% NP-40) supplemented with cOmplete™ Protease Inhibitor Cocktail (Sigma), PhosSTOP™ (Roche), and 1 mM PMSF. After sonication and centrifugation (13,000 g, 30 min), the supernatant was incubated with FLAG antibody-conjugated Dynabeads™ Protein G (Invitrogen) at 4□ for 3 h, washed five times with IP-1000 buffer, and eluted with 3x FLAG Peptide (Protein Ark). Plasmids are listed in Key resources table.

### In vitro binding assay

For in vitro binding of GST-RIOK2-His, RIOK2 and CLK1 recombinant proteins were incubated with in vitro kinase assay buffer (50 mM Tris-HCl pH 7.5, 150 mM NaCl, 5 mM MgCl_2_, 2 mM DTT) at 37□ for 30 min. Proteins were suspended in IP-500 buffer (50 mM Tris-HCl pH 7.5, 500 mM NaCl, 0.5% NP-40) supplemented with 1 mM PMSF and mixed with Glutathione Sepharose™ 4 Fast Flow (Cytiva). After rotating at 4□ for 2 h, proteins were washed five times and eluted with SDS sample buffer. For in vitro binding of MBP-FLAG-PPP1CC-His, PPP1CC and proteins of interest were incubated with NEBuffer for PMP (NEB) at 37□ for 30 min. Protein complexes were purified using Amylose Resin (NEB).

To assess the temperature-dependence of the interaction between RIOK2 or SRSF9 and CLK1, GS4B-bound RIOK2 or SRSF9 was mixed with CLK1 in IP-150 buffer containing 5 mM MgCl_2_ and incubated at 37□ or 42□ for 30 min with shaking at 950 rpm. After washing the complexes five times with IP-150 buffer preheated to 37□ or 42□, the complexes were eluted using SDS sample buffer.

To assess the temperature-dependent dissociation of PPP1R2, PPP1R8, or CLK1 from PPP1CC, complexes were first formed by incubation at 37□ for 30 min and purified using Amylose Resin (NEB) at 4□. The purified complexes were then divided into two and incubated at either 37□ or 42□ for 15 min. After collecting the supernatant, the resin-bound complexes were washed with preheated NEBuffer for PMP (NEB) and eluted with SDS sample buffer.

To evaluate the binding affinity between RIOK2 and dephosphorylated CLK1, recombinant CLK1 was incubated with Amylose Resin (NEB) bound either MBP-FLAG-PPP1CC-His or MBP-FLAG-His in NEBuffer for PMP (NEB) supplemented with 1 mM PMSF and 1mM MnCl_2_ at 37□ for 30 min. Following incubation, the NaCl concentration was raised to 500 mM to disrupt non-specific interactions, and the resin-bound MBP-tagged proteins were separated from the supernatant. The supernatant was then mixed with GST-RIOK2-His recombinant protein, and the solution was diluted with IP-0 buffer (50 mM Tris-HCl pH 7.5, 0.5% NP-40) to adjust the NaCl concentration to 150 mM. GST pulldown was subsequently performed using Glutathione Sepharose™ 4 Fast Flow resin (Cytiva). Bound complexes were eluted with SDS sample buffer and analyzed by western blotting.

### Isothermal titration calorimetry (ITC)

For ITC experiments, His-tag purified PPP1CC and PPP1R2 recombinant proteins were desalted in dialysis buffer using Superdex™ 200 Increase 10/300 GL and HiTrap™ Desalting columns (Cytiva), respectively. Measurements were performed using a Microcal PEAQ-ITC calorimeter (Malvern Panalytical) at 750 rpm and either 37□ or 42□. The sample cell and syringe contained 23.5 μM PPP1CC and 162 μM PPP1R2, respectively. Titration consisted of an initial 0.4 μl injection followed by 18 injections of 2 μl at 150-second intervals. Data were analyzed with MicroCal PEAQ-ITC software (Version 1.41). The first injection was excluded from analysis at both 37□ and 42□, and the second injection at 42□ was also excluded as an outlier. As a control, PPP1R2 was injected into dialysis buffer alone, confirming that only constant dilution heat was detected. Thermodynamic parameters—enthalpy change (ΔH), dissociation constant (Kd), and binding stoichiometry (N)—were determined by fitting to a single-site binding model. In the analysis of the 42□ data, the N value was set to the same value determined from the 37□ data. Errors represent fitting uncertainties.

### In vitro kinase assay

RIOK2 and CLK1 recombinant proteins were incubated with in vitro kinase assay buffer containing 0.1-0.2 MBq [γ-^32^P] labeled ATP (Revvity) at 37□ for 30 min. The phosphorylation levels were analyzed by SDS-PAGE and autoradiography using a BAS-SR imaging plate (Fujifilm) and scanned with a Typhoon FLA 9500 scanner (GE Healthcare Life Sciences). CQ211 was added simultaneously with the onset of incubation. In parallel, recombinant proteins were incubated under the same conditions with cold ATP (Cytoskeleton, Inc.) instead of [γ-³²P]-labeled ATP to validate protein levels by western blotting.

### In vitro phosphatase assay

PPP1CC and either PPP1R2 or PPP1R8 recombinant proteins were incubated with NEBuffer for PMP (NEB) on ice for 30 min. CLK1 recombinant protein in NEBuffer for PMP supplemented with 1 mM PMSF and 1mM MnCl2 was then mixed, and samples were incubated at 37□ or 42□ for 40 min. After incubation, MnCl2 was removed by TCA precipitation, and samples were analyzed by Phos-tagTM SDS-PAGE and western blotting.

### In vitro reconstitution of CLK1 phosphorylation status

For the in vitro reconstitution system to assess how PP1 and RIOK2 regulate CLK1 phosphorylation at 37□ and 42□, recombinant PPP1CC and PPP1R2 proteins were pre-incubated on ice for 30 min. Subsequently, RIOK2, CLK1, and the PPP1CC–PPP1R2 complex were incubated in kinase assay buffer containing 0.1–0.2 MBq [γ-^32^P] ATP (Revvity) and 1 mM MnCl at 37□ or 42□ for 30 min. Total protein amounts were normalized by supplementing with BSA. Following incubation, samples were subjected to SDS-PAGE, and radiolabeled proteins were detected by autoradiography as described above.

To reconstitute the temperature changes during recovery from thermal stress, recombinant RIOK2, CLK1, PPP1CC, and PPP1R2 were incubated in kinase assay buffer at 42□ for 30 min. After this preincubation, 0.1–0.2 MBq [γ-32P] ATP (Revvity) was added, and the reactions were either shifted to 37□ or maintained at 42□ for 30 min. Samples were subjected to SDS-PAGE and autoradiography as described above.

Protein levels in these assays were validated using the same methods as described for the in vitro kinase assay (see above).

## QUANTIFICATION AND STATISTICAL ANALYSIS

qRT-PCR results were quantified using LightCycler 480 software (version 1.5.1, Roche). Western blot, FISH and IF images were analyzed with ImageJ (FIJI). Pearson’s R value was calculated using Coloc 2 in ImageJ. Statistical significance was assessed using GraphPad Prism 10. P-values are indicated by asterisks: ****p < 0.0001, ***0.0001 < p < 0.001, **0.001 < p < 0.01, *0.01 < p < 0.05.

## REFERENCES

1. Richter, K., Haslbeck, M., and Buchner, J. (2010). The Heat Shock Response: Life on the Verge of Death. Mol Cell 40, 253–266.

2. Guil, S., and Cáceres, J.F. (2007). Stressful Splicing. Mol Cell 28, 180–181.

3. Ninomiya, K., Kataoka, N., and Hagiwara, M. (2011). Stress-responsive maturation of Clk1/4 pre-mRNAs promotes phosphorylation of SR splicing factor. J Cell Biol 195, 27–40.

4. Chanseok Shin, Ying Feng, and James L. Manley (2004). Dephosphorylated SRp38 acts as a splicing repressor in response to heat shock. Nature 427, 548–553.

5. Shalgi, R., Hurt, J.A., Lindquist, S., and Burge, C.B. (2014). Widespread inhibition of posttranscriptional splicing shapes the cellular transcriptome following heat shock. Cell Rep 7, 1362–1370.

6. Isabel Novoa, Y.Z.H.Z.R.J.H.P.H. and D.R. (2003). Stress induced gene expression requires programmed recovery from translational repression. EMBO J 22, 1180–1187.

7. Kim, D., Ouyang, H., and Li, G.C. (1995). Heat shock protein hsp70 accelerates the recovery of heat-shocked mammalian cells through its modulation of heat shock transcription factor HSF1. Proc Natl Acad Sci USA 92, 2126–2130.

8. Biamonti, G., and Vourc’h, C. (2010). Nuclear stress bodies. Cold Spring Harb Perspect Biol 2, a000695.

9. Riggs, C.L., Kedersha, N., Ivanov, P., and Anderson, P. (2020). Mammalian stress granules and P bodies at a glance. J Cell Sci 133.

10. Anderson, P., and Kedersha, N. (2008). Stress granules: the Tao of RNA triage. Trends Biochem Sci 33, 141–150.

11. Molliex, A., Temirov, J., Lee, J., Coughlin, M., Kanagaraj, A.P., Kim, H.J., Mittag, T., and Taylor, J.P. (2015). Phase Separation by Low Complexity Domains Promotes Stress Granule Assembly and Drives Pathological Fibrillization. Cell 163, 123–133.

12. Yang, P., Mathieu, C., Kolaitis, R.M., Zhang, P., Messing, J., Yurtsever, U., Yang, Z., Wu, J., Li, Y., Pan, Q., et al. (2020). G3BP1 Is a Tunable Switch that Triggers Phase Separation to Assemble Stress Granules. Cell 181, 325–345.e28.

13. Yuan, L., Mao, L.H., Huang, Y.Y., Outeiro, T.F., Li, W., Vieira, T.C.R.G., and Li, J.Y. (2025). Stress granules: emerging players in neurodegenerative diseases. Transl Neurodegener 14, 22.

14. Chujo, T., Yamazaki, T., Kawaguchi, T., Kurosaka, S., Takumi, T., Nakagawa, S., and Hirose, T. (2017). Unusual semi-extractability as a hallmark of nuclear body-associated architectural noncoding RNAs. EMBO J 36, 1447–1462.

15. Hirose, T., Ninomiya, K., Nakagawa, S., and Yamazaki, T. (2023). A guide to membraneless organelles and their various roles in gene regulation. Nat Rev Mol Cell Biol 24, 288–304.

16. Fujiwara, N., Ueno, T., Yamazaki, T., and Hirose, T. (2025). Unraveling architectural RNAs: Structural and functional blueprints of membraneless organelles and strategies for genome-scale identification. Biochim Biophys Acta Gen Subj 1869, 130815.

17. Hirose, T., Fujiwara, N., Ninomiya, K., Yamamoto, T., Nakagawa, S., and Yamazaki, T. (2025). Architectural RNAs: blueprints for functional membraneless organelle assembly. Trends Genet.

18. Altemose, N., Logsdon, G.A., Bzikadze, A. V., Sidhwani, P., Langley, S.A., Caldas, G. V., Hoyt, S.J., Uralsky, L., Ryabov, F.D., Shew, C.J., et al. (2022). Complete genomic and epigenetic maps of human centromeres. Science 0376, eabl4178.

19. Rhie, A., Nurk, S., Cechova, M., Hoyt, S.J., Taylor, D.J., Altemose, N., Hook, P.W., Koren, S., Rautiainen, M., Alexandrov, I.A., et al. (2023). The complete sequence of a human Y chromosome. Nature 621, 344–354.

20. Denegri, M., Moralli, D., Rocchi, M., Biggiogera, M., Raimondi, E., Cobianchi, F., De Carli, L., Riva, S., and Biamonti, G. (2002). Human chromosomes 9, 12, and 15 contain the nucleation sites of stress-induced nuclear bodies. Mol Biol Cell *13*, 2069–2079.

21. Jolly, C., Konecny, L., Grady, D.L., Kutskova, Y.A., Cotto, J.J., Morimoto, R.I., and Vourc’h, C. (2002). In vivo binding of active heat shock transcription factor 1 to human chromosome 9 heterochromatin during stress. Journal of Cell Biology 156, 775–781.

22. Ninomiya, K., Adachi, S., Natsume, T., Iwakiri, J., Terai, G., Asai, K., and Hirose, T. (2020). Lnc RNA -dependent nuclear stress bodies promote intron retention through SR protein phosphorylation. EMBO J 39, e102729.

23. Ninomiya, K., Yamazaki, T., and Hirose, T. (2023). Satellite RNAs□: emerging players in subnuclear architecture and gene regulation. EMBO J 42, e114331.

24. Jolly, C., Metz, A., Govin, J., Vigneron, M., Turner, B.M., Khochbin, S., and Vourc’h, C. (2004). Stress-induced transcription of satellite III repeats. J Cell Biol 164, 25–33.

25. Rizzi, N., Denegri, M., Chiodi, I., Corioni, M., Valgardsdottir, R., Cobianchi, F., Riva, S., and Biamonti, G. (2004). Transcriptional Activation of a Constitutive Heterochromatic Domain of the Human Genome in Response to Heat Shock. Mol Biol Cell 15, 543–551.

26. Liu, X.Q., Li, P., Gao, B.Q., Zhu, H. Le, Yang, L.Z., Wang, Y., Zhang, Y.Y., Wu, H., Pan, Y.H., Shan, L., et al. (2025). De novo assembly of nuclear stress bodies rearranges and enhances NFIL3 to restrain acute inflammatory responses. Cell. 188, 4586–4603.e31.

27. Denegri, M., Chiodi, I., Corioni, M., Cobianchi, F., Riva, S., and Biamonti, G. (2001). Stress-induced Nuclear Bodies Are Sites of Accumulation of Pre-mRNA Processing Factors. Mol Biol Cell 12, 3502–3514.

28. Metz, A., Soret, J., Vourc’h, C., Tazi, J., and Jolly, C. (2004). A key role for stress-induced satellite III transcripts in the relocalization of splicing factors into nuclear stress granules. J Cell Sci 117, 4551–4558.

29. Mähl, P., Lutz, Y., Puvion, E., and Fuchs, J.-P. (1989). Rapid Effect of Heat Shock on Two Heterogeneous Nuclear Ribonucleoprotein-associated Antigens in HeLa Cells. J Cell Biol 109, 1921–1935.

30. Sarge, K.D., Murphy, S.P., and Morimoto, R.I. (1993). Activation of Heat Shock Gene Transcription by Heat Shock Factor 1 Involves Oligomerization, Acquisition of DNA-Binding Activity, and Nuclear Localization and Can Occur in the Absence of Stress. Mol Cell Biol 13, 1392–1407.

31. Ninomiya, K., Iwakiri, J., Aly, M.K., Sakaguchi, Y., Adachi, S., Natsume, T., Terai, G., Asai, K., Suzuki, T., and Hirose, T. (2021). m^6^A modification of HSATIII lncRNAs regulates temperature-dependent splicing. EMBO J 40, e107976.

32. Shin, Y., and Brangwynne, C.P. (2017). Liquid phase condensation in cell physiology and disease. Science 357, eaaf4382.

33. Shi, Y., and Manley, J.L. (2007). A Complex Signaling Pathway Regulates SRp38 Phosphorylation and Pre-mRNA Splicing in Response to Heat Shock. Mol Cell 28, 79–90.

34. Boutz, P.L., Bhutkar, A., and Sharp, P.A. (2015). Detained introns are a novel, widespread class of post-transcriptionally spliced introns. Genes Dev 29, 63–80.

35. Jiang, X., Liu, B., Nie, Z., Duan, L., Xiong, Q., Jin, Z., Yang, C., and Chen, Y. (2021). The role of m^6^A modification in the biological functions and diseases. Signal Transduct Target Ther 6:74.

36. Cerezo, E.L., Houles, T., Lie, O., Sarthou, M.K., Audoynaud, C., Lavoie, G., Halladjian, M., Cantaloube, S., Froment, C., Burlet-Schiltz, O., et al. (2021). RIOK2 phosphorylation by RSK promotes synthesis of the human small ribosomal subunit. PLoS Genet 17.

37. LaRonde-LeBlanc, N., and Wlodawer, A. (2005). The RIO kinases: An atypical protein kinase family required for ribosome biogenesis and cell cycle progression. Biochimica et Biophysica Acta - Proteins and Proteomics, pp. 14–24.

38. Ghosh, S., Raundhal, M., Myers, S.A., Carr, S.A., Chen, X., Petsko, G.A., and Glimcher, L.H. (2022). Identification of RIOK2 as a master regulator of human blood cell development. Nat Immunol 23, 109–121.

39. LaRonde-LeBlanc, N., and Wlodawer, A. (2005). A family portrait of the RIO kinases. J Biol Chem 280, 37297–37300.

40. Zemp, I., Wild, T., O’Donohue, M.F., Wandrey, F., Widmann, B., Gleizes, P.E., and Kutay, U. (2009). Distinct cytoplasmic maturation steps of 40S ribosomal subunit precursors require hRio2. Journal of Cell Biology 185, 1167–1180.

41. Ouyang, Y., Si, H., Zhu, C., Zhong, L., Ma, H., Li, Z., Xiong, H., Liu, T., Liu, Z., Zhang, Z., et al. (2022). Discovery of 8-(6-Methoxypyridin-3-yl)-1-(4-(piperazin-1-yl)-3-(trifluoromethyl)phenyl)-1,5-dihydro-4H-[1,2,3]triazolo[4,5- c]quinolin-4-one (CQ211) as a Highly Potent and Selective RIOK2 Inhibitor. J Med Chem *65*, 7833–7842.

42. Mohamed, A.A., Xavier, C.P., Sukumar, G., Tan, S.H., Ravindranath, L., Seraj, N., Kumar, V., Sreenath, T., McLeod, D.G., Petrovics, G., et al. (2018). Identification of a small molecule that selectively inhibits ERG-positive cancer cell growth. Cancer Res 78, 3659–3671.

43. Fedorov, O., Huber, K., Eisenreich, A., Filippakopoulos, P., King, O., Bullock, A.N., Szklarczyk, D., Jensen, L.J., Fabbro, D., Trappe, J., et al. (2011). Specific CLK inhibitors from a novel chemotype for regulation of alternative splicing. Chem Biol 18, 67–76.

44. Kim, Y., Holland, A.J., Lan, W., and Cleveland, D.W. (2010). Aurora kinases and protein phosphatase 1 mediate chromosome congression through regulation of CENP-E. Cell 142, 444–455.

45. Xin, G., Fu, J., Luo, J., Deng, Z., Jiang, Q., and Zhang, C. (2020). Aurora B regulates PP1g-Repo-Man interactions to maintain the chromosome condensation state. J Biol Chem 295, 14780–14788.

46. Heroes, E., Lesage, B., Görnemann, J., Beullens, M., Van Meervelt, L., and Bollen, M. (2013). The PP1 binding code: A molecular-lego strategy that governs specificity. FEBS J 280, 584–595.

47. Foulkes, J.G., and Cohen, P. (1980). The Regulation of Glycogen Metabolism: Purification and Properties of Protein Phosphatase Inhibitor-2 from Rabbit Skeletal Muscle. Eur J Biochem 105, 195–203.

48. Tsuboyama, K., Osaki, T., Matsuura-Suzuki, E., Kozuka-Hata, H., Okada, Y., Oyama, M., Ikeuchi, Y., Iwasaki, S., and Tomari, Y. (2020). A widespread family of heat-resistant obscure (Hero) proteins protect against protein instability and aggregation. PLoS Biol 18.

49. Cho, N.H., Cheveralls, K.C., Brunner, A.D., Kim, K., Michaelis, A.C., Raghavan, P., Kobayashi, H., Savy, L., Li, J.Y., Canaj, H., et al. (2022). OpenCell: Endogenous tagging for the cartography of human cellular organization. Science (1979) 375.

50. Haltenhof, T., Kotte, A., De Bortoli, F., Schiefer, S., Meinke, S., Emmerichs, A.K., Petermann, K.K., Timmermann, B., Imhof, P., Franz, A., et al. (2020). A Conserved Kinase-Based Body-Temperature Sensor Globally Controls Alternative Splicing and Gene Expression. Mol Cell 78, 57–69.e4.

51. Taniguchi, I. (2024). The SARS-CoV-2 ORF6 protein inhibits nuclear export of mRNA and spliceosomal U snRNA. PLoS One 19.

