## Supplementary figures and images for "Thermo-Sensing Mechanisms of Splicing Control by Nuclear Stress Bodies"

### Supplemental Figures S1-S6

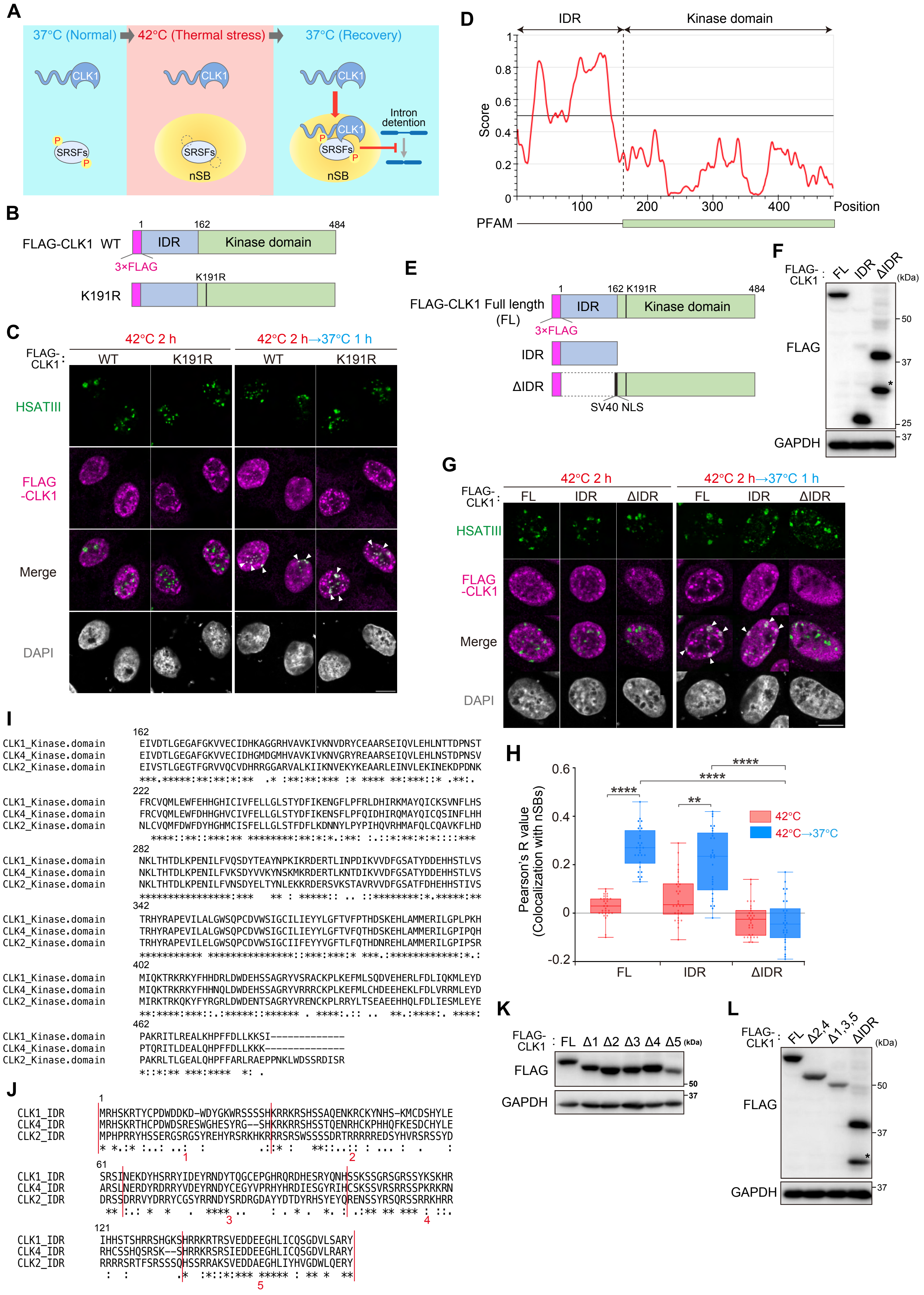

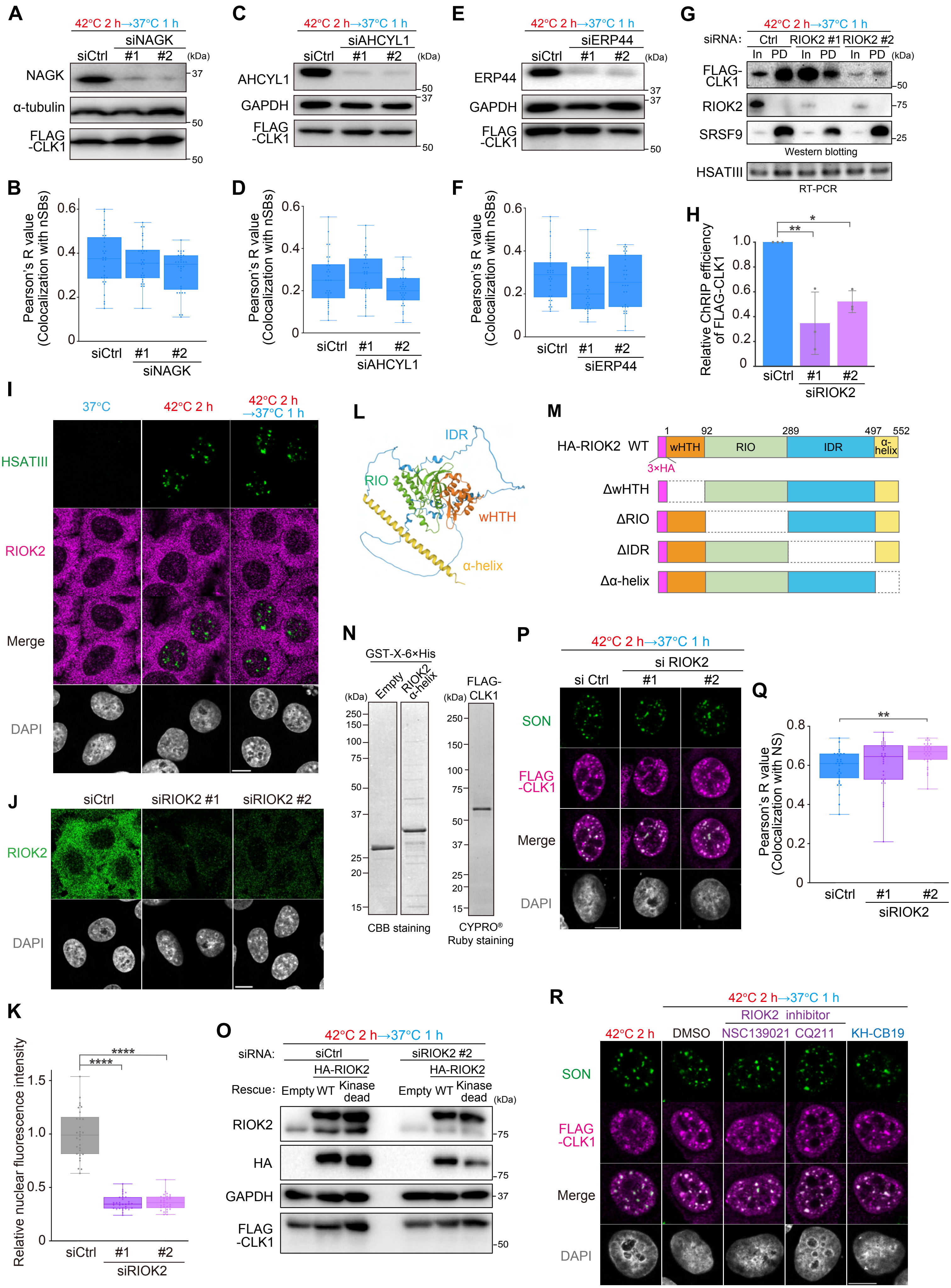

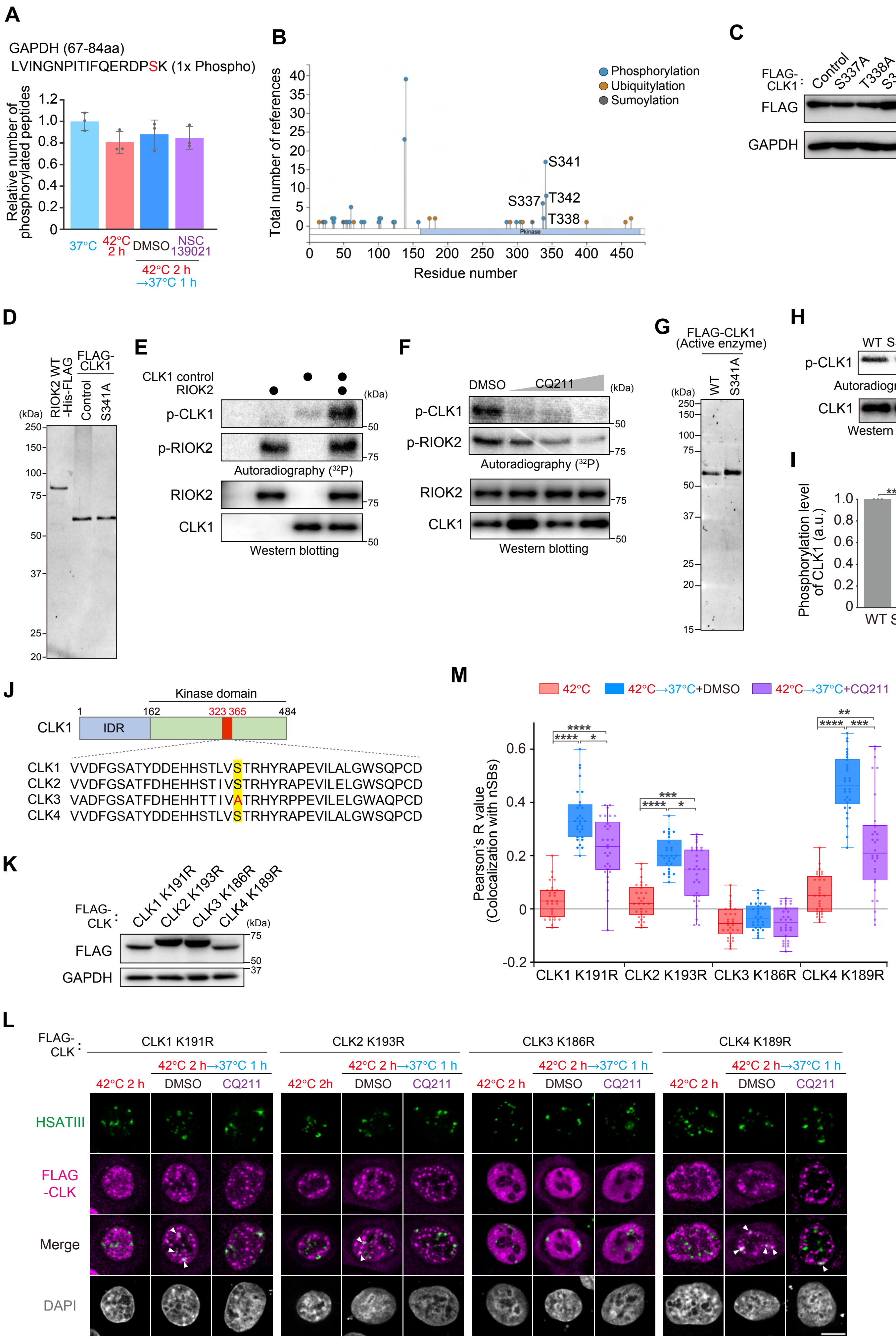

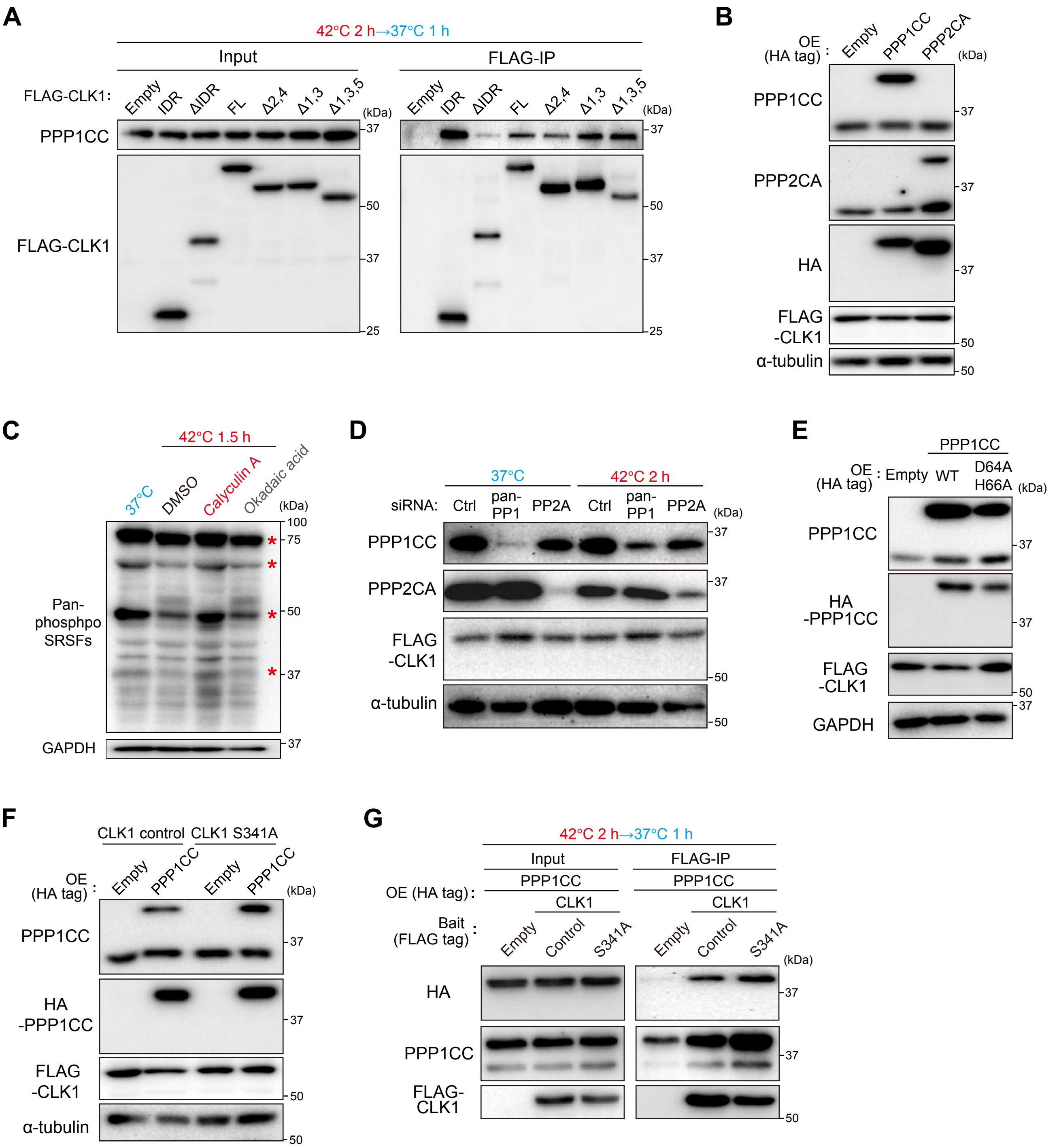

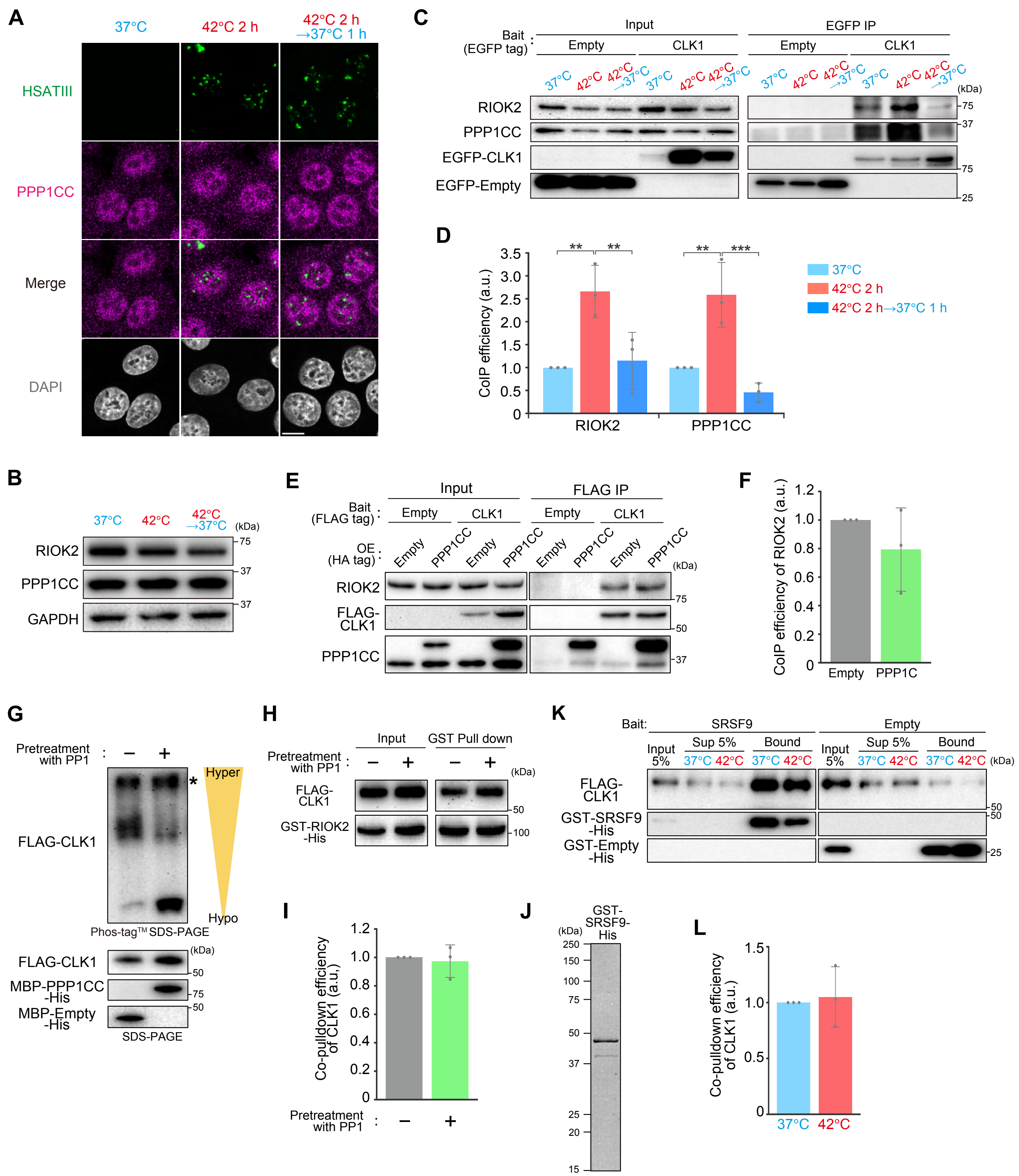

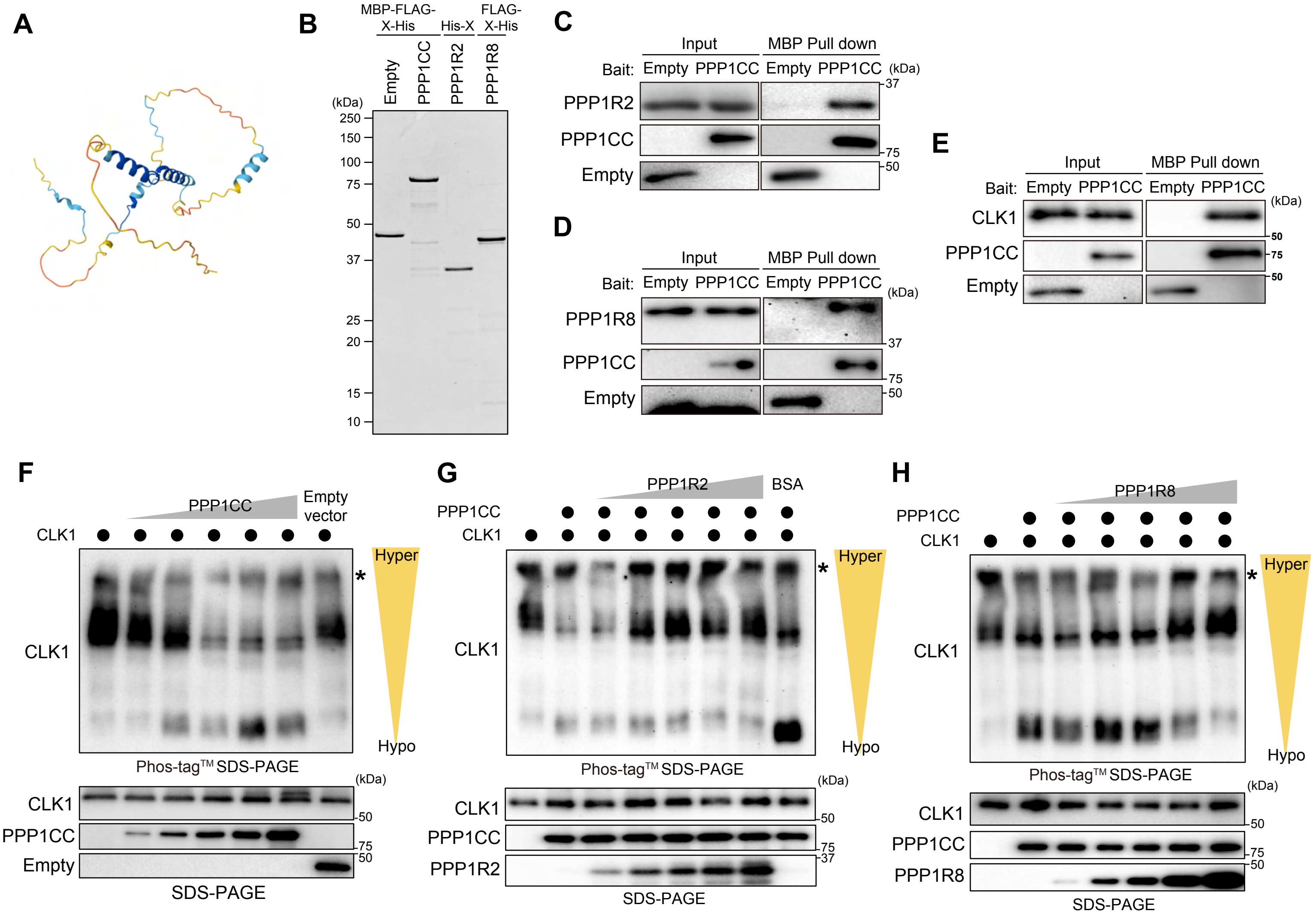
