## Supplemental Figure legends for "Thermo-Sensing Mechanisms of Splicing Control by Nuclear Stress Bodies"

**Figure S1. Identification of essential regions within CLK1’s IDR for temperature-dependent localization to nSBs, related to Figure 1**

(A) Model for temperature-dependent CLK1 localization and reaction crucible function of nSBs in regulating temperature-dependent pre-mRNA splicing.

(B, E) Schematic of domain composition of FLAG-CLK1 and its mutants. (B) K191R, a kinase-dead mutant. (E) Full length (FL) and ΔIDR constructs carry the K191R mutation.
(C, G) Subnuclear localization of CLK1 WT and mutants under thermal stress and recovery, visualized by HSATIII RNA-FISH combined with FLAG IF. White arrowheads indicate HSATIII-CLK1 colocalization. Nuclei were counterstained with DAPI. Scale bar, 10 μm.
(D) Prediction of CLK1 IDR using IUPred3.
(F, K, L) Expression of CLK1 mutants validated by western blotting (WB). Asterisks indicate a partial fragment of the ΔIDR mutant (F, L).
(H) Quantification of colocalization of CLK1 variants with nSBs (n = 30) in (G). *****p* < 0.0001, **0.001 < *p* <0.01 (Dunn’s multiple comparisons test).
(I, J) Amino acid sequence alignments of CLK1, CLK2, and CLK4. (H) Kinase domain. (I) N-terminal IDR. Number represents amino acid positions of CLK1. Alignment was performed using CLUSTALW.

**Figure S2. Screening for regulator proteins of CLK1 localization to nSBs and characterization of RIOK2, related to Figure 2**

(A, C, E) Evaluation of knockdown efficiency of candidate regulators by WB. α-tubulin and GAPDH are loading controls.
(B, D, F) Quantification of colocalization of CLK1 with nSBs upon knockdown of the regulator candidates in (A, C, E). Box plots show Pearson’s correlation coefficients (R values) between HSATIII-FISH and FLAG-IF signals in the nucleus (n = 30). Dunn’s multiple comparisons test was used.
(G) HSATIII-ChIRP during thermal stress recovery upon RIOK2 knockdown, followed by WB. Protein input (In) 2%. HSATIII lncRNA was detected by RT-PCR, with 25% of the total RNA input loaded (In). PD, pulldown products of HSATIII-ChIRP.

(H) Quantification of ChIRP efficiency of FLAG-CLK1 in (G). ChIRP efficiency was determined by calculating the ratio of PD to In. Data are presented as mean ± SD (n = 3). **0.001 < *p* <0.01, *0.01 < *p* < 0.05 (Dunnet’s multiple comparisons test).
(I) Subcellular localization of RIOK2 across temperature conditions, visualized by IF combined with HSATIII RNA-FISH. Scale bar, 10 μm.
(J) Validation of RIOK2 IF signal specificity using RIOK2 siRNAs.
(K) Quantification of nuclear fluorescence intensities of RIOK2 signals in (J) (n = 30). *****p* < 0.0001 (Dunn’s multiple comparisons test).

(L) Structural prediction of RIOK2 generated using AlphaFold2.

(M) Schematic of domain composition of HA-RIOK2 WT and its mutants.

(N) Detection of purified recombinant WT or mutant RIOK2 and CLK1 proteins by SDS-PAGE and CYPRO^®^ Ruby staining.
(O) Evaluation of RIOK2 knockdown efficiency and the expression level of RIOK2 rescue plasmid, assessed by WB.
(P, R) Subnuclear localization of CLK1 during thermal stress and recovery, visualized by SON IF (green) combined with FLAG IF (magenta). (P) Cells were transfected with siRNAs (20 nM). (R) DMSO or inhibitors (CQ211, 10 μM; NSC139021, 30 μM) were added during recovery Scale bar, 10 μm.
(Q) Quantification of colocalization of CLK1 with nuclear speckles upon RIOK2 knockdown in (P) (n = 30). **0.001 < *p* <0.01 (Dunn’s multiple comparisons test).

**Figure S3. Identification of the phosphorylation site within CLK1 by RIOK2 and its contribution to CLK1 localization to nSBs, related to Figure 3**

(A) Bar graph showing relative abundance of phosphorylation of GAPDH at Ser83 under the indicated condition. Data are presented as mean ± SD (n = 3), with each sample measured twice by mass spectrometry and averaged. Šidák’s multiple comparisons test was used.

(B) Phosphorylation sites within CLK1 listed in the PhosphoSitePlus® database.
(C, K) WB of (C) FLAG-CLK1 amino acid substitution mutants and (K) FLAG-CLK1-4.
(D) Detection of purified recombinant RIOK2 and CLK1 proteins by SDS-PAGE and CYPRO^®^ Ruby staining.
(E, F) In vitro assessment of RIOK2-mediated phosphorylation of CLK1 and RIOK2 itself by autoradiography with western blot validation. (F) CQ211 was added at 10, 20, and 50 μM.
(G) Detection of purified catalytically active recombinant CLK1 WT and S341A mutant by SDS-PAGE and CYPRO^®^ Ruby staining.

(H) In vitro assessment of CLK1 autophosphorylation by autoradiography with western blot validation.
(I) Quantification of autoradiographic signals of CLK1 in (H). Data are presented as mean ± SD (n = 3). ***0.0001 < *p* < 0.001 (two-tailed t-test).
(J) Schematic of the aligned amino acid sequence of CLK1 (323–365 aa) and corresponding regions in other CLK family proteins.
(L) Subnuclear localization of kinase-dead mutants of CLK family proteins under thermal stress and recovery conditions, visualized by HSATIII RNA-FISH combined with FLAG IF. DMSO or CQ211 (10 μM) was added during recovery. Scale bar, 10 μm.
(M) Quantification of colocalization of CLK mutants with nSBs in (L) (n = 30). *****p* < 0.0001, ***0.0001 < *p* < 0.001, **0.001 < *p* <0.01, *0.01 < *p* < 0.05 (Dunn’s multiple comparisons test).

**Figure S4. Identification of the phosphatase responsible for dephosphorylation of CLK1 during thermal stress, related to Figure 4**

(A) Detection of CLK1-PPP1CC interaction by coIP from cell lysates during stress recovery, followed by WB. FLAG-CLK1 variants used for coIP are indicated above the panels.
(B, E, F) Evaluation of expression levels of transfected PPP1CC WT, PPP1CC mutant or PPP2CA by WB. α-tubulin and GAPDH are loading controls.
(C) Detection of pan-phosphorylated SRSFs using the 1H4 antibody. Asterisks indicate bands dephosphorylated during thermal stress. GAPDH is a loading control.
(D) Evaluation of PP1 and PP2A knockdown efficiency by WB. α-tubulin is a loading control.
(G) Detection of PPP1CC-CLK1 control/S341A by coIP from cell lysates during stress recovery, followed by WB.

**Figure S5. Temperature-dependent interaction between CLK1 and RIOK2, related to Figure 5**

(A) Subcellular localization of PPP1CC across temperature conditions, visualized by IF combined with HSATIII RNA-FISH. Scale bar, 10 μm.
(B) Evaluation of RIOK2 and PPP1CC expression across temperature conditions by WB,

(C) CLK1 interactions with RIOK2 and PPP1CC across temperature conditions by coIP　of EGFP-CLK1 followed by WB to detect RIOK2 and PPP1CC.
(D) Quantification of coIP efficiency of RIOK2 and PPP1CC in (C). CoIP efficiency was calculated as the ratio of EGFP IP band intensity of RIOK2/PPP1CC to that of CLK1. Data are presented as mean ± SD (n = 3). ***0.0001 < *p* < 0.001, **0.001 < *p* <0.01 (Šidák’s multiple comparisons test).
(E) Detection of CLK1–RIOK2 interactions by coIP from cell lysates under normal temperature, upon overexpression of empty vector or PPP1CC, followed by WB.
(F) CoIP efficiency of RIOK2 in (E). CoIP efficiency was calculated as the ratio of FLAG IP band intensity of RIOK2 to that of CLK1. Data are presented as mean ± SD (n = 3). Two-tailed t-test was used.

(G) Monitoring of the phosphorylation status of FLAG-CLK1 pretreated with recombinant MBP-PPP1CC or MBP alone (empty vector control) by Phos-tag™ SDS-PAGE (upper panel) and normal SDS-PAGE (lower panel) followed by WB. Migration of different phosphorylation states is shown on the right. Asterisks indicate putative aggregated CLK1 proteins.
(H) Direct interactions between recombinant CLK1 and RIOK2 proteins pretreated with recombinant MBP-PPP1CC or MBP alone by GST pulldown and WB.
(I) Quantification of co-pulldown efficiency of CLK1 in (H). Co-pulldown efficiency was calculated as the ratio of the band intensity of CLK1 to that of RIOK2 in the GST pull-down fraction. Data are presented as mean ± SD (n = 3). Two-tailed t-test was used.
(J) Detection of purified recombinant SRSF9 proteins by SDS-PAGE and CBB staining.
(K) Detection of direct interactions between recombinant CLK1 and SRSF9 at 37℃ or 42℃ by GST pull-down and WB.
(L) Quantification of co-pulldown efficiency of CLK1 in (K). Co-pulldown efficiency was calculated as the ratio of the band intensity of CLK1 to that of SRSF9 in the GST pulldown fraction. Data are presented as mean ± SD (n = 3). Two-tailed t-test was used.

**Figure S6. Thermo-sensing mechanisms of PP1, related to Figure 6**

(A) Structural prediction of PPP1R2 generated using AlphaFold2.
(B) Detection of purified recombinant PPP1CC, PPP1R2 and PPP1R8 proteins by SDS-PAGE and CBB staining. Protein tags are indicated above the panel.
(C–E) Direct interactions between recombinant MBP-PPP1CC and PPP1R2/PPP1R8/CLK1 proteins by MBP pulldown and WB.
(F–H) Monitoring of CLK1 phosphorylation status after incubation with: (F) PPP1CC or empty vector; (G) PPP1CC combined with PPP1R2 or BSA; (H) PPP1CC combined with PPP1R8. Phosphorylation states were assessed by Phos-tag™ SDS-PAGE followed by WB. Asterisks indicate putative aggregated CLK1 proteins.
